# Coordinated Response Modulations Enable Flexible Use of Visual Information

**DOI:** 10.1101/2024.07.10.602774

**Authors:** Ramanujan Srinath, Martyna M. Czarnik, Marlene R. Cohen

## Abstract

We use sensory information in remarkably flexible ways. We can generalize by ignoring task-irrelevant features, report different features of a stimulus, and use different actions to report a perceptual judgment. These forms of flexible behavior are associated with small modulations of the responses of sensory neurons. While the existence of these response modulations is indisputable, efforts to understand their function have been largely relegated to theory, where they have been posited to change information coding or enable downstream neurons to read out different visual and cognitive information using flexible weights. Here, we tested these ideas using a rich, flexible behavioral paradigm, multi-neuron, multi-area recordings in primary visual cortex (V1) and mid-level visual area V4. We discovered that those response modulations in V4 (but not V1) contain the ingredients necessary to enable flexible behavior, but not via those previously hypothesized mechanisms. Instead, we demonstrated that these response modulations are precisely coordinated across the population such that downstream neurons have ready access to the correct information to flexibly guide behavior without making changes to information coding or synapses. Our results suggest a novel computational role for task-dependent response modulations: they enable flexible behavior by changing the information that gets out of a sensory area, not by changing information coding within it.

**Significance:** Natural perceptual judgments are continuous, generalized, and flexible. We estimate the ripeness of a piece of fruit on a continuous scale, we generalize by judging the ripeness of either a mango or an avocado even though they look very different, we flexibly judge either the size or the ripeness of the same piece of fruit, and we can flexibly indicate the same perceptual judgment using a variety of behaviors such as by speaking or writing any of many languages. Here, we show that the response modulations in visual cortex long associated with cognitive processes, surround modulation, or motor planning are sufficient to guide all these aspects of natural perceptual decision-making. We find that across the population, these response modulations reorient and reformat visual representations so that the relevant information is used to guide behavior via communication with downstream neurons. Our results are an example of a general computational principle for flexible behavior that emerges from the coordinated activity of large populations of neurons.

## Introduction

Perceptual, cognitive, and motor processes have long been known to modulate the responses of sensory neurons. In visual cortex, processes including contrast, adaptation, surround modulation, attention, learning, task switching, arousal, action planning and more are associated with modest modulations of neural responses(*1–7*). These modulations are broadly consistent with multiplicatively scaling (changing the gain of) relatively stable tuning curves(*8*). More recently, modulation related to a broad set of perceptual, cognitive, and motor processes have been observed in essentially every brain area in many species(*3*, *9*, *10*).

Two related frameworks have dominated thinking about the function of these signals. The first is the idea that these modulations change the information *encoded* in the population because high gains are associated with improved signal-to-noise ratio(*11–13*). The second idea is that mixing perceptual, cognitive, and motor signals (e.g. ‘mixed selectivity’) benefits the flexible use of those signals(*14*, *15*) because in theory, representations of different perceptual, cognitive, and motor features could be flexibly *decoded* from neural activity, typically using a different weighted combination of neural responses for each variable(*14*, *16–18*).

We tested a third possibility, that response modulations in visual cortex coordinate across the population to change how visual features are represented so that the appropriate information is communicated via a fixed weighted combination of projection neurons. Put another way, we hypothesize that many aspects of population responses are fixed, including the information encoded in a population and the weighted combination of neurons that communicates with downstream areas to guide behavior. Behavioral flexibility comes from transforming those representations (e.g. by rotating them) such that the information communicated by that fixed combination of neurons is flexible and task dependent. Using coordinated response modulations as a substrate for behavioral flexibility is consistent from what we know about the biology and function of neural circuits. It gives downstream neurons access to the relevant subset of information without requiring synaptic changes or other circuit level changes that might be costly and time consuming. The notion of fixed output neurons is consistent with anatomical observations that a stable subset of neurons projects from any area to each downstream target (*19*).

To test this hypothesis, we recorded groups of neurons in visual areas V1 and V4 while monkeys performed a perceptual estimation task that allows us to strongly link neural responses to choices on a continuous scale (Figure 1A-B). Our results demonstrate that the tuning and coordinated, task- and stimulus-dependent modulation of responses of populations of neurons in V4 (but not in V1) contain all the ingredients to support flexible perceptual decision-making. Both V1 and V4 neurons robustly encode both task-relevant and irrelevant visual features and a stable linear combination of those responses predicts the monkeys’ perceptual judgments across stimuli and task conditions. We show that V4 neurons are modulated in a coordinated way such that the stable combination of neurons reflects the direction of the upcoming eye movement or which of two visual features would be discriminated. We also demonstrate that these coordinated gain changes are consistent with, but do not necessarily follow from, the results of past studies that focus on single neurons.

**Figure 1:**
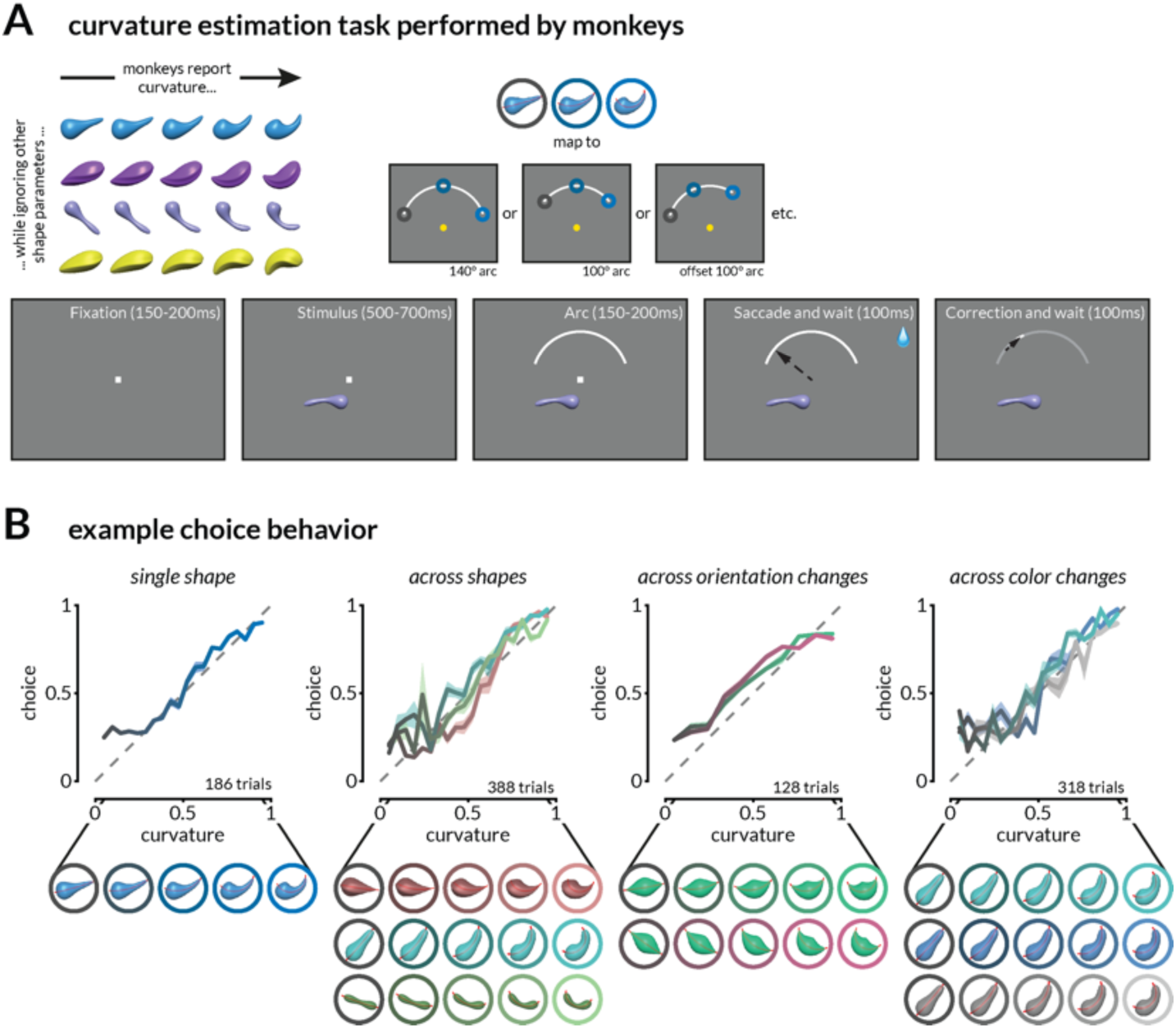
Task and behavior. **A.** Schematic of the continuous curvature estimation task. Stimuli that varied in curvature and task-irrelevant features were presented in the joint receptive fields of V1 and V4 neurons as monkeys fixated a central dot. After 550-800ms, a target arc was presented in the upper hemifield, and monkeys were rewarded for making a saccade to a location on the arc that corresponded to the stimulus curvature. The reward amount was inversely related (with a threshold) to the error in curvature judgment. In a subset of experiments, the radial position, angular position, and length of the target arc were varied pseudorandomly. **B.** Monkeys report medial axis curvature while ignoring other stimulus features. Example continuous estimation behavior on four sessions during which the curvature of one or many shapes was estimated on interleaved trials. Shading indicates the standard error of the mean (SEM).

## Results

Gain changes related to attention(*1*), task-switching(*2*), arousal(*3*), visual context(*4*, *5*), and motor planning(*6*) have several characteristics: they are typically modest in size(*8*), they depend on the relationship between the neuron’s tuning and the specific modulatory process (e.g. the focus of attention, the surrounding stimulus, or the planned action), and they are heterogeneous, meaning that even simultaneously recorded neurons with indistinguishable tuning can show different amounts of modulation(*20*, *21*). This heterogeneity means that single neuron responses are insufficient to infer the impact of modulation on the population representation.

Depending on how those gain changes are coordinated across the population, those modulations could change information coding by changing the signal-to-noise ratio of neural responses(*1*, *22*), average out and have minimal impact(*23*), or, as we propose here, coordinate to fundamentally change the information used to guide behavior via a fixed combination of readout neurons(*24*, *25*).

To investigate the relationship between these gain changes and flexible behavior, we measured stimulus and task-related responses of populations of neurons at multiple stages of visual processing. We designed a behavioral framework that required monkeys to exhibit many forms of flexibility: they made continuous perceptual judgments, generalized across stimuli with many task-irrelevant features to make a single judgment, mapped that single judgment onto one of multiple actions, and switched which visual feature was task-relevant. We trained two rhesus monkeys to perform a continuous curvature estimation task (Figure 1A-B) while we recorded neural activity from areas V1 and V4 using chronically implanted multi-electrode arrays (Figure S3A). After fixating on a central spot, a stimulus was presented in the joint receptive fields of the recorded neurons (which was always in the lower hemifield; Figure S3B). After the monkeys viewed the stimulus, they indicated their judgment about the curvature by looking at a corresponding position on an arc presented in the upper hemifield. A saccade to the leftmost portion of the arc indicated a straight stimulus (curvature = 0), a saccade to the right rightmost end corresponded to the maximum curve presented to the animals (curvature = 1), and the monkey could indicate any intermediate value in between. The monkeys were rewarded according to the accuracy of their estimate (see Methods). We generated the stimuli by randomizing the cross-sectional shape, aspect ratio, color, 3D rotation, twist along the axis, gloss, thickness profile, and other features of an oblong object (Figure 1A and S1). On any given session, stimuli were drawn from three to seven base shapes that differed in features such as overall shape, color, or orientation, each with 10-20 curvatures. The monkeys learned to report the curvature of the presented objects while ignoring irrelevant feature changes (Figure 1B and S2). During training sessions, monkeys were exposed to 10-15 randomly generated shapes which were repeated during recording sessions with additional changes in color and orientation. In a subset of sessions, we also presented random shapes that the monkeys had never seen before. Curvature judgement error patterns between familiar and novel shapes were similar for both monkeys (Figure S2C and S2D). Additionally, in some sessions, we changed the angular position and length of the target arc (randomly interleaved from trial to trial), so the monkeys select their specific eye movement after seeing the arc (they were rewarded for choosing the correct relative position on the arc, which varied according to the position and length of the arc on that trial).

V1 and V4 neurons responded to the curvature, color, and orientation of the stimuli (Figure S3C). The curvature selectivity of neurons was shape-dependent and did not vary systematically with the onset or details of the arc (Figure S3D-E). While V4 neurons have been shown to support coding transformations that extract curvature(*26–28*) and V1 neurons to extract 2D curvature(*29*, *30*), several image-level changes are correlated with curvature in these shapes. We use ‘curvature’ representation as a shorthand for the representation of the changing feature that the monkey is trained to report and not the explicit representation of medial axis curvature.

### A stable weighted combination of V1 and V4 neurons explains perceptual judgments amid many task-irrelevant stimulus changes

A key prediction of our hypothesis is that although task-irrelevant visual features might greatly affect the responses of visual neurons, the representations must enable the animal to use a general strategy (such as basing choices on a stable linear combination of neural responses) to make a perceptual judgment for any stimulus. We tested this prediction by assessing the relationship between the V1 and V4 representations of the curvature of different shapes and the animals’ curvature judgment.

Both V1 and V4 contain representations of curvature that are strikingly different for different shapes. In both areas, population responses to shapes that vary from straight to curvy trace out smooth, uninterrupted paths in neural population space where each dimension is the firing rate of one neuron (Figure 2B, S4). We can estimate the curvature of each shape by finding the linear combination of neurons that best accounts for curvature (this traces out an axis in neural population space), and we quantify the “shape-specific” curvature information as the correlation between the predicted and actual curvature on held out trials (Figure 2C).

**Figure 2:**
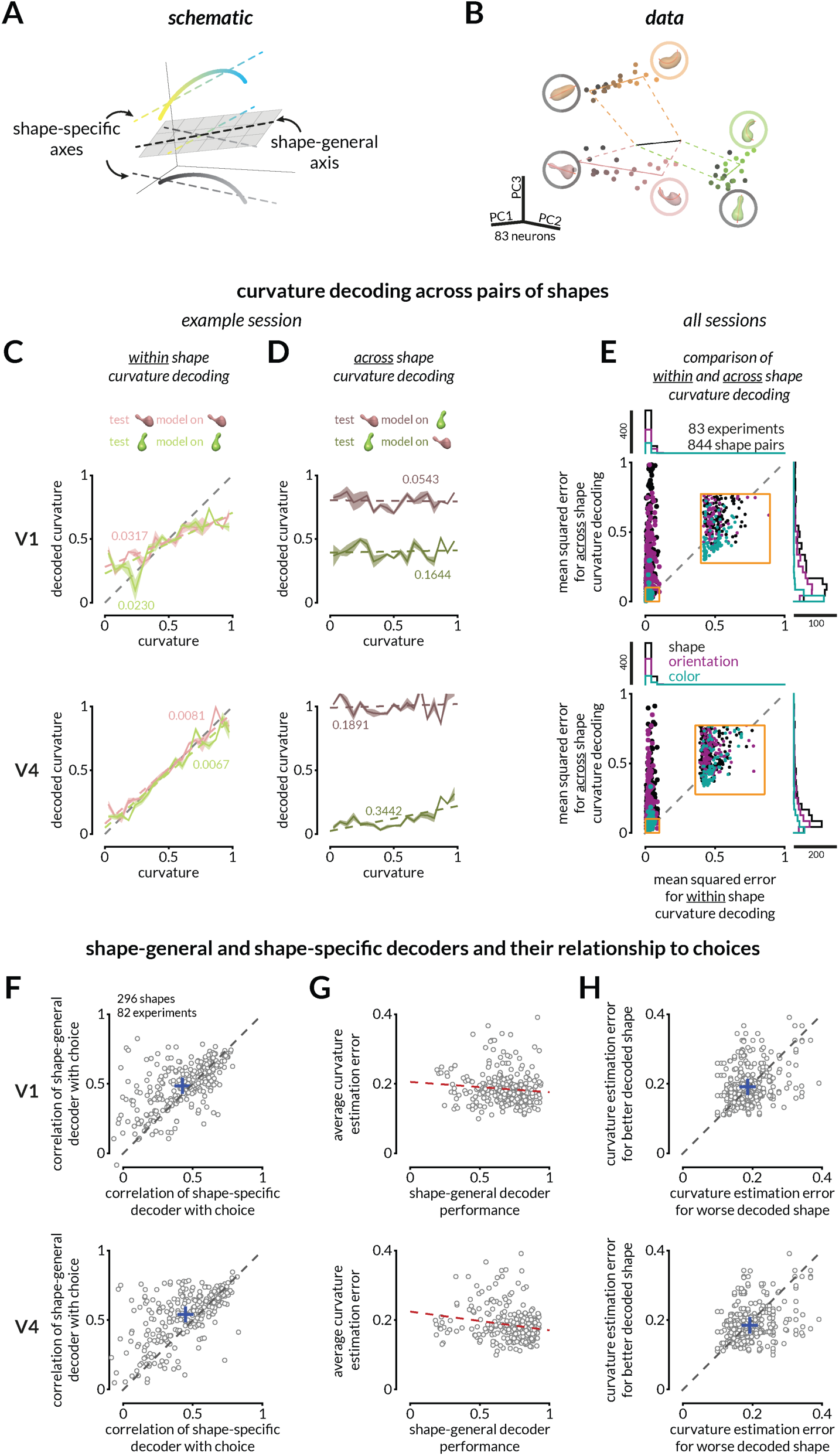
The monkeys’ behavior suggests that a fixed linear combination of neural responses explains curvature judgments across many irrelevant stimulus changes. **A.** The representations of curvature for different shapes can be misaligned but a common shape-general axis can represent curvature. **B.** For illustration purposes, V4 population responses to three shapes (varying in curvature) recorded on interleaved trials are projected into the first three principal components of the space in which the response of each simultaneously recorded neuron is one dimension. The points indicate the average population response to each unique stimulus at each curvature value. Luminance of the points (black to pink, green, or orange) represents increasing curvature. Solid colored lines represent best fit lines for each shape (shape-specific axis), and the black line represents the shape-general axis (see Figure S4 for other examples). **C.** Curvature can be linearly decoded (cross-validated) from V1 (top) and V4 (bottom) responses for each shape independently for the example session in B. Colored labels indicate mean squared error (MSE) between the actual and predicted curvatures, and colored lines depict the linear fit relating predicted and actual curvatures. Shading indicates SEM across 100 folds of 50% trial splits (see Figure S4 for more example decoding analyses.) **D.** The representations of curvature are not aligned across shapes. Same decoders as C, but the linear decoder is trained on responses to one shape and tested on responses to another in V1 (top) or V4 (bottom). **E.** Curvature representations for different shapes are typically misaligned. Across shape pairs, our ability to decode curvature from V1 (top) and V4 (bottom) is better when the weights are based on responses to the same shape (x-axis) than when it is based on responses to the other shape in the pair (y-axis); MSE for same shape decoding from V1 responses (left) and V4 responses (right) is smaller than for across shapes (Wilcoxon rank sum test; p < 10^-10^). Each dot indicates a shape pair that is different in overall shape appearance (black), orientation only (purple), or color only (teal). The inset is a zoomed in view of the indicated area on the plot. The marginal distributions are shown on the top and right. **F.** The monkeys’ choices are more strongly correlated with the prediction of a shape-general linear combination of neural responses (as in C) than a shape-specific strategy for V1 (top) and V4 (bottom) (Wilcoxon signed rank tests; p<10^-8^ V1; p<10^-14^ V4). Each point represents the correlation values (between the decoded curvature and the monkey’s choice) for one shape. **G.** Behavioral choices are more accurate (lower average absolute error) for shapes whose representation is well-aligned with the shape-general curvature decoder of V4 (bottom) responses (correlation r= –0.17, vs constant model p=0.003), but not significantly for V1 (correlation r= –0.07, vs constant model p=0.261). **H.** For pairs of shapes tested in the same session, average behavioral error was typically bigger for the worse decoded shape (using a shape-general decoder; x-axis) than the better decoded shape (y-axis) in V4 (Wilcoxon signed rank test; V4: p=0.004), but not significantly for V1 (p=0.19).

Changing the shape (Figure S4B), orientation (Figure S4C), or color (Figure S4D) of the stimulus resulted in misaligned shape-specific representations, meaning that different weights best decoded the curvature of shapes with different task-irrelevant features. To quantify this misalignment, we attempted to estimate the curvature of one shape using the weighted linear combination of neurons that best predicted curvature of a different shape that was recorded on randomly interleaved trials (Figure 2). For 100 cross-validation folds, we calculated the weights for one shape using half of the trials and tested it either on the other half (“within shape curvature decoding” – Figure 2C) or on the responses to a different shape (“across shape curvature decoding” – Figure 2D). Even though the monkey’s curvature estimation behavior across the two shapes was consistently shape-independent (Figure S5A), across shape decoding was poor in both V1 and V4 (Figure 2E). Across sessions, the behavioral estimation of curvature across pairs of stimuli that varied in overall shape, orientation, or color was highly correlated (Figure S5B), but the across-shape curvature decoding consistently underperformed within-shape curvature decoding (Figure 2E).

We reasoned that the monkeys might employ a shape-general strategy, meaning that they use a common linear combination of neural responses to estimate the curvature of any stimulus. A shape-general strategy generates three testable predictions: (1) the monkeys’ choices should be better correlated with predictions of the shape-general decoder than the shape-specific decoder, (2) monkeys should be better at estimating the curvature of shapes whose representations happen to be better aligned with the shape-general axis, and (3) between any two shapes, the shape whose representation is better aligned with the shape-general axis should have lower behavioral estimation errors on average. We tested these predictions by training a shape-general decoder on V1 or V4 responses to all of the shapes presented in a given session. Consistent with our hypothesis, the monkeys’ choices were better correlated with curvature decoded along the shape-general than shape-specific axes in both V1 (Figure 2F top) and V4 (bottom). This result was consistent across sessions in which the shape-general decoder was trained on objects that varied in overall shape, only orientation or only color (Figure S6). In addition, shapes with higher average estimation errors by the monkeys were also poorly decoded by the shape-general decoder (Figure 2G), and between pairs of shapes that were tested on randomly interleaved trials, the shape for which curvature more poorly aligned with the shape-general axis was more poorly estimated by the monkeys (higher average behavioral errors; Figure 2H). Together, these results demonstrate that both V1 and V4 contain stimulus representations that are sufficient to enable shape-general curvature judgments.

### Response modulations in V4, but not V1, reformat curvature representations to enable flexible mapping of one perceptual judgment to many actions

In our task, monkeys can make a perceptual judgment as soon as the stimulus is displayed but must wait to map that judgment to an action (eye movement) until the arc is displayed. We dissociated the curvature judgment from the eye movement by varying the angular position and length of the target arc across trials. We posited that coordinated gain changes associated with the arc onset (attributable to surround modulation, drawing attention to the arc, or a combination) are coordinated in a way that transforms the curvature representation in a way that could guide the appropriate eye movement. To test the viability of this idea, we simulated neurons with curvature tuning functions whose gain is modulated slightly by the presentation of two different arcs (Figure 3A-B). We chose modest, heterogeneous gains for each neuron by drawing from the same distribution for each of two simulated arc conditions. Some random draws produced gains that would transform the curvature representation so that a fixed linear combination of simulated neurons would communicate the appropriate eye movement for that arc (Figure 3B, dark and light green). However, other draws from the same distribution did not transform the population response appropriately, meaning that the specified linear combination of neurons did not produce appropriate actions (Figure 3B, gray). This simulation demonstrates that the modulations in single neurons observed in previous studies are consistent with, but do not imply, the idea that they coordinate across the population to transform population responses to enable an appropriate mapping between a perceptual judgment and the action used to communicate it.

**Figure 3:**
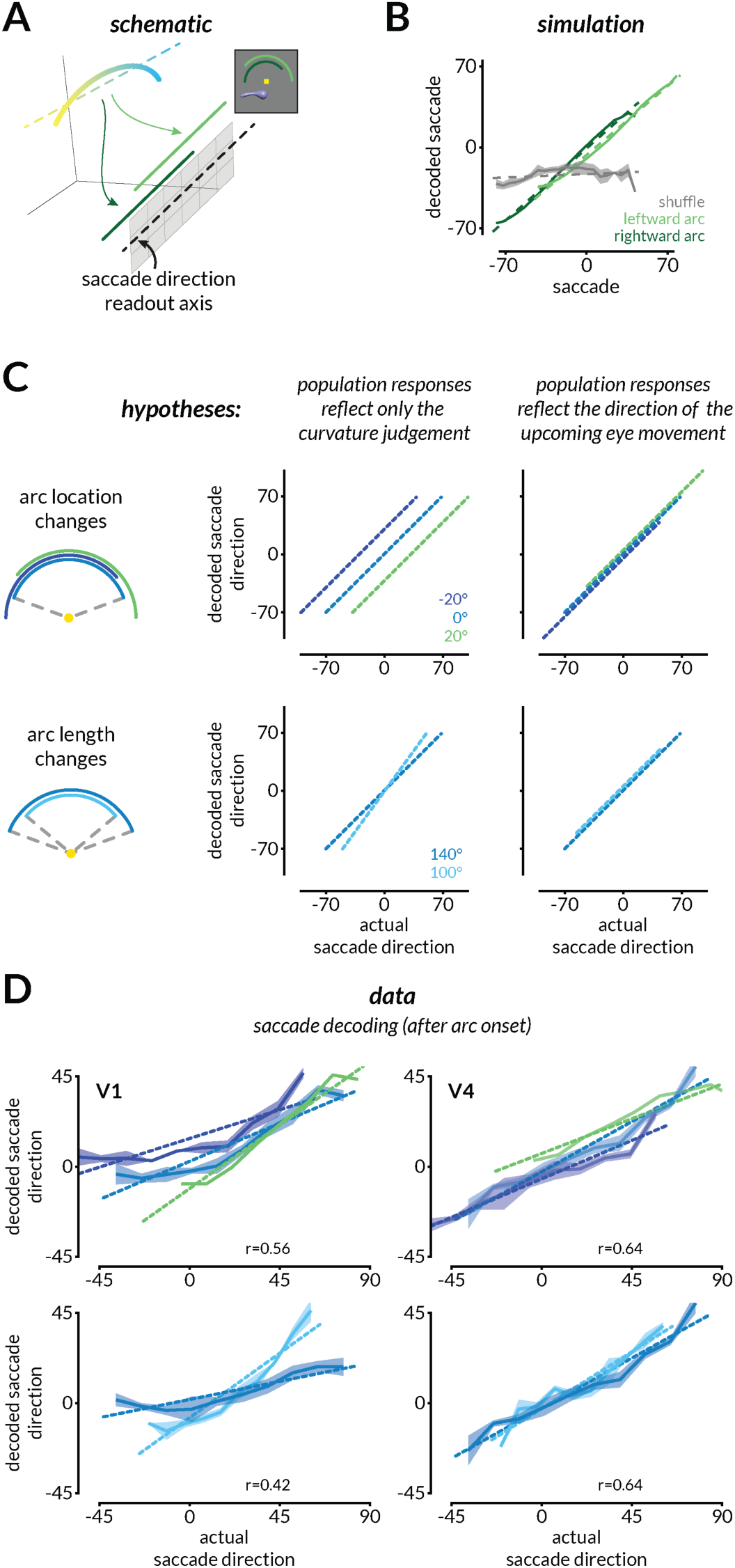
V4, but not V1, population activity is reformatted to enable the flexible mapping of curvature to different actions. **A.** To enable a flexible mapping between curvature judgments and eye movement responses, the curvature representation of a shape could be transformed to align with a fixed readout axis that communicates with eye movement planning areas. In other words, the same curvature judgment should be mapped to different parts of the saccade direction axis depending on the location and length of the arc. **B.** In simulations, we randomly assigned the arc-dependent gain modulations to neurons by drawing from a distribution of response gains that is consistent with single neuron results (see Methods). Only some draws of the same distribution enabled the mapping of curvature representations to the appropriate portions of the readout axis such that the saccade could be decoded from the population (light and dark green). Most other draws did not (gray). **C.** Schematic depicting analyses that would reveal whether neural population responses reflect only the curvature judgment and not the upcoming eye movement used to communicate that judgment (left) or whether they also reflect the upcoming eye movement (right). Colors represent predictions for the different arc locations (top) or lengths (bottom). **D.** V4, but not V1, responses reflect the direction of the upcoming eye movement in example sessions. The impending saccade direction was decoded from V1 (left) and V4 (right) responses during the period when the monkeys have not yet moved their eyes but after the onset of the arc that allows them to plan the eye movement. Shading indicates SEM and the correlation between actual and predicted saccade direction is labeled on the bottom-right of each panel.

To test this hypothesis, we compared the neural representations before and after the target arc was displayed. Since the monkey was required to maintain fixation for a short interval after the onset of the arc and before making the saccade, we posited that a modest gain change (which might be attributable to any combination of surround modulation, attention, action planning, or other modulatory processes) should transform the curvature representation such that we should be able to linearly decode the impending saccade direction from population responses. This is somewhat counterintuitive for visual cortex, which might be expected to largely encode the curvature judgment without specific premotor signals (although see (*31*, *32*) about premotor signals in V4). In the absence of premotor signals, the decoded saccade would not depend on the location or length of the arc (Figure 3C left).

We found that V4, but not V1, representations were transformed to encode the direction of the upcoming eye movement (Figure 3D). Both areas represented the curvature judgment before the onset of the arc (Figure 2 and S4). After the onset of the arc, the small gain changes in V4 transformed the representation so that it encoded the direction of the upcoming eye movement (Figure 3D, right and S7C). In contrast, V1 continued to represent the curvature judgment (Figure 3D left and S7B).

V4 is one of many areas in the monkey brain that encode the direction of an upcoming eye movement. To test the hypothesis that transforming a representation from a curvature judgment to an action plan could be a plausible substrate for this sort of flexible behavior, we created a model that has only one brain area. We trained a recurrent neural network (RNN) to map a stimulus curvature to a saccade given an arc condition with the same trial structure (Figure S8). Like the V4 neurons, the RNN hidden units represented the stimulus curvature and then, after arc onset, transformed the representation based on the arc condition to align with an axis encoding the direction of the upcoming saccade (Figure S8E). This model demonstrates that transforming the representation in an arc-dependent manner is a viable computational substrate for mapping one perceptual judgment onto many actions. We also extended this model by adding a pre-trained convolutional neural network (CNN) front-end that supplied the RNN its “visual inputs” in the form of hidden layer activations (Figure S9A). Like V1 and V4 neural populations, the RNN learned a shape-general curvature axis from CNN activations and transformed the representation to align with a saccade direction axis (Figure S9B). Our results demonstrate that small gain changes in visual cortex are sufficient to enable behavioral generalization by mapping many visual features to one judgment (Figure 2), and behavioral flexibility by the mapping of one judgment onto multiple actions (Figure 3).

### Response modulations enable a fixed combination of V4 neurons to reflect choices about curvature or color

The strongest test of our hypothesis that we could think of is the case when choices can be based on either of two visual features (e.g. curvature and color) that are represented in the same group of neurons. In this scenario, gain changes consistent with task switching, feature attention, and/or surround modulation must rotate the population representation so that information about the relevant feature is communicated to decision neurons. We therefore extended our hypotheses (Figure 4A) and simulations (Figure 4B) to incorporate neural tuning to two features and corresponding gain changes consistent with the single neuron modulations observed in studies that require task-switching or feature attention. These studies find that while there is substantial neuron-to-neuron heterogeneity, neurons tend to have higher gains when monkeys perform a task for which they are better tuned(*22*). We therefore simulated tuning functions (Figure S11A) to two features and assigned each neuron small selectivity-dependent gains (Figure 4B).

**Figure 4:**
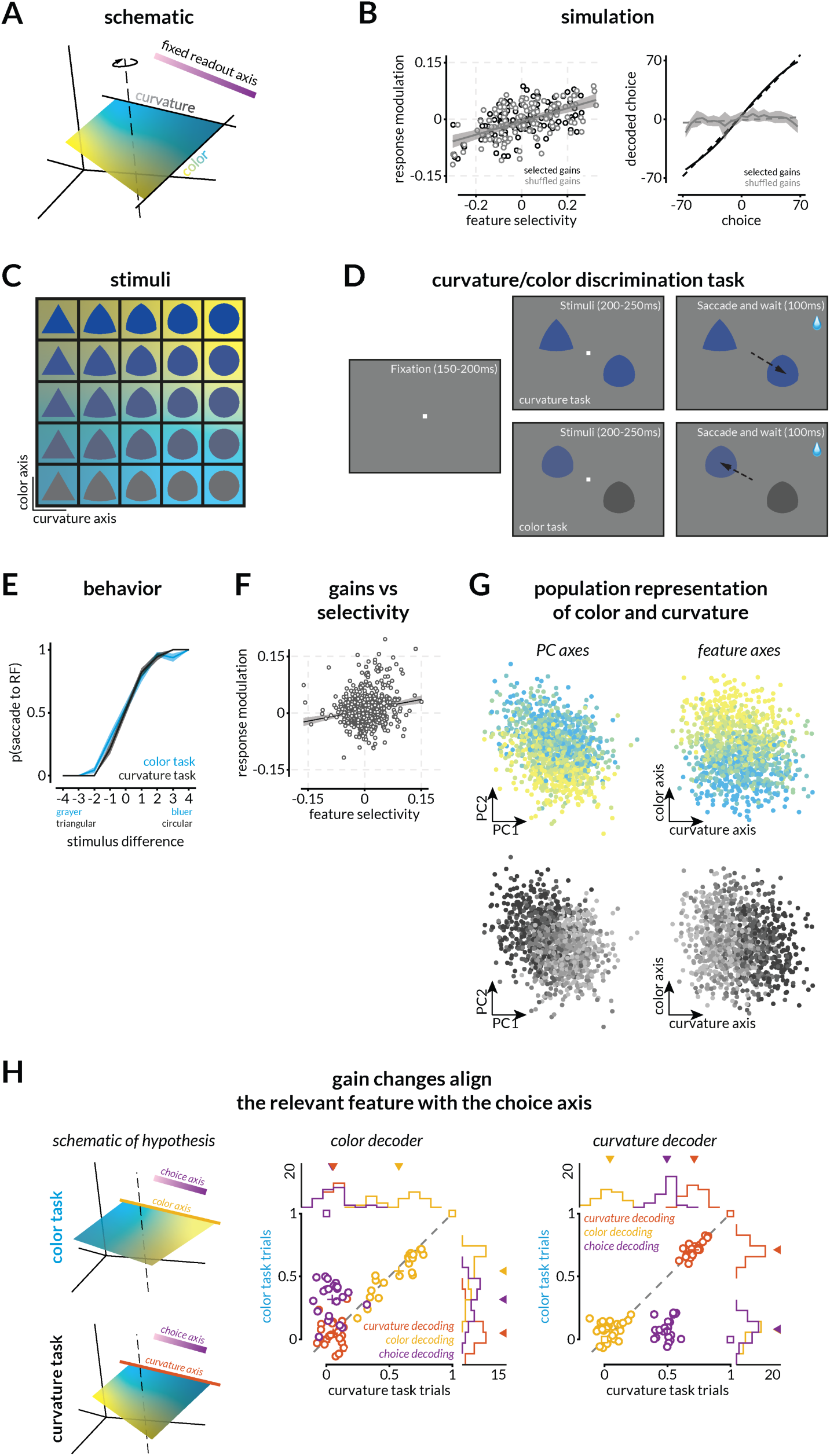
Small gain changes consistent with feature attention or surround modulation enable different visual features to guide choices. **A.** Schematic of the hypothesis that gain changes will reformat stimulus representations to align with a fixed readout axis depending upon the relevant feature. **B.** Small gain changes can, but do not necessarily transform the population response to enable different features to guide choices. Similar to Figure 3, we simulated neurons tuned to two features and assigned small gains to each neuron by drawing from a distribution similar to the distribution of gains that have been reported for feature attention(*22*). Some random draws transform the population such that the representation of the task-relevant feature is aligned with the readout axis (black). As in the previous simulation, many random draws from the same distributions (black vs gray points; left) do not (gray; right). **C.** Stimuli varied in color/luminance (blue to gray) and curviness/shape (triangle to circle). Background colors allow comparison to neural results in G and H. **D.** Schematic of the two-feature (curvature/color) discrimination task. The monkey was rewarded for making a saccade to one of the stimuli (one in the joint receptive fields of the recorded V4 neurons and one in the opposite hemifield). During the curvature task, the colors of the two stimuli were the same (selected from the same row in C) and the more circular stimulus was rewarded. During the color task, the stimuli were the same (selected from the same column in C) and the bluer stimulus was rewarded. **E.** Example psychometric curves for the curvature (black) and color (blue) task. The plot depicts the proportion of trials in which the monkey chose the stimulus in the receptive fields of the recorded V4 neurons as a function of the relevant feature of the stimulus that was in that receptive field. These data are from a single experimental session (314 total trials, 89% correct overall; 176 color task trials, 89% correct; 138 shape task trials, 90% correct). Across the 23 sessions that were used for further analysis, during which curvature and color task trials were randomly interleaved, the monkey performed at 85.45% correct overall (∼ 610 trials on average per session), 79.54% on the color task, and 91.65% on the shape task. **F.** Replication of previous results showing a relationship between the modulation of neural responses associated with the different tasks and the selectivity of the neuron to the two features. The linear fit (R^2^=0.032; intercept=0.007 (p<10^-11^); slope=0.19 (p<10^-8^)) and 95% confidence intervals are indicated by black line and gray shaded region. **G.** Population representation of the two stimulus features using PCA (left) and QR decomposition (right) to visualize the representations of color (blue to yellow gradient) and shape (black to gray gradient) of the image in the population RF. Each point is the V4 population response on one trial. **H.** Evidence that task-dependent gains reformat neural representations to enable flexibility in which feature guides choices (left). The middle and right plots depict correlations between the values predicted by the linear combination of V4 responses that best predicts color (left) or curvature (right) and the actual values of shapes, colors, and choices in the color (y-axes) or curvature task trials (x-axes). Across both tasks, shape and color are decoded well on the shape and color axes respectively, but not vice versa, suggesting that the two features are approximately orthogonally represented in neural population space. As predicted by the simulations (square markers), on color task trials, we could decode the animal’s choices significantly better on the color than on the shape axis (Wilcoxon signed rank test, p<10^-4^), and on shape trials, we could decode choices significantly better on the shape than the color axis (Wilcoxon signed rank test, p<10^-4^). Marginal histograms with arrows indicating means are shown at the top and right of both panels (black line and the number indicate the experiment count).

Our simulations demonstrated that these small gain changes can, but do not necessarily, transform the neural representations to enable different features to guide choices using a fixed, task-general decision axis. We found that some random draws from the same distribution of gain changes (Figure 4B, left) transformed the population such that we could decode the value of the task-relevant feature on a fixed decision axis (Figure 4B, right, black line) while many draws did not (gray line).

We tested this hypothesis experimentally using a simplified two-alternative forced choice task where either the color (gray to blue) or the curviness (triangle to circle) of the displayed shape (Figure 4C) predicted the correct answer. The monkey was rewarded for selecting the bluer stimulus (color task) or the more circular one (curvature task; Figure 4D-E). We placed one stimulus in the joint receptive fields of recorded V4 neurons (Figure 4D and S10B) and the other in the opposite hemifield. The recorded neurons had a range of tuning preferences (Figure S10C) and displayed similar patterns and magnitudes of task-dependent gain modulation as previously published (Figure 4F; (*22*)). The V4 populations we recorded contained smooth and uninterrupted population representations of both shape and color (Figure 4G). Our hypothesis and simulations predict that the monkey’s choices will be based on, and therefore decodable from, the representation of the task-relevant visual feature, but not the irrelevant one (Figure 4H; left). We tested this prediction by identifying the axes in neural space that best encode shape and color and then determining whether we could decode choices from projections along those axes. There was plenty of information about both the relevant and irrelevant stimulus features when the monkey performed both the shape and color tasks, but choices were only aligned with the task-relevant feature (Figures 4H, middle and right, and Figure S12B-C). These results demonstrate that task- or stimulus-specific modulations can transform visual representations to enable flexible discrimination of any visual feature to which the neurons are tuned.

## Discussion

Together, our results show that long-known response modulations of V4 neurons coordinate in precise ways that would allow them enable flexible visually guided behavior. We demonstrated four ways that V4 populations are well suited to enable flexible behavior:

1) The representations of the relevant feature for each shape (curvature), while strongly modulated by task-irrelevant features, were smooth and continuous (Figure 2 and S4) and therefore well positioned to mediate continuous perceptual judgments.
2) The monkeys’ perceptual judgments were most closely associated with a fixed, shape-general strategy (Figure 2, S5, and S6), which is well positioned to enable the rapid estimates of even novel stimuli in novel scenes that are characteristic of natural behavior.
3) Modulations at the onset of the target arc transformed the neural representations in a way that could map a single judgment to many possible actions via a fixed combination of neurons (Figure 3 and S7).
4) Modulations associated with different task-relevant stimulus features (shape vs. color) transformed the population representation such that neural information about the task-relevant feature could be communicated via a fixed combination of neurons (Figure 4 and S12).

These results provide broad support for the idea that well-known, small modulations of visual neurons provide a computational substrate for cognitive and behavioral flexibility via relatively fixed communication or readout strategies. These results suggest the tantalizing possibility that the reformatting of neural population representations could be a brain-wide substrate for flexible behavior.

### Flexible reformatting of population representations is consistent with observations about the stability of the tuning of single neurons and with known biological processes

Across different tasks, cognitive states, stimuli, and motor plans, neurons in visual cortex largely retain their tuning to visual features and undergo modest task-specific modulations(*1*). Because of this tuning stability, cognitive flexibility is typically attributed to flexible interactions between neural populations(*25*, *33*). The implied neural mechanism passes off the computational burden of inferring the task-relevant stimulus features and mapping to behavioral choices to the modulation or gating of synapses between different neurons that communicate with each other via dedicated channels or communication subspaces.

Here, we tested the alternative, but not mutually exclusive, hypothesis that flexibility comes from reformatting neural representations such a fixed subset of readout neurons could selectively reflect the perceptual, cognitive, and motor information necessary to guide behavior. Only a subset of neurons in any brain area project to a given downstream target(*19*), which is consistent with the idea that the readout strategy (e.g. by downstream decision neurons) is largely stable. Attempts to identify the subset of information that is functionally shared between any two neural populations (i.e. the ‘communication subspace’(*33*)) have shown that those interactions are largely stable(*34*), even though the information shared via that stable subspace can vary(*25*). While our results suggest that relevant information is identified in visual cortex and communicated to downstream areas to guide decisions, they do not preclude channels for additionally communicating task-irrelevant information to downstream decision areas (*35*). Together, these results and those presented here are consistent with the idea that at least for forms of behavioral flexibility that can change on the timescale of a single trial, the output channels (defined loosely as projection neurons, communication subspaces, and/or readout strategies) are largely fixed. An interesting question for future work is whether flexible processes that evolve on longer timescales, such as perceptual or motor learning, might involve synaptic changes that modify the output channels.

### Role of distinct brain areas in flexible behavior

If modulations to visual neurons contain all the ingredients for flexible, visually guided behavior, why does the monkey have so many other brain areas? An emerging body of evidence, including that presented here, is that the formatting of different information sources (e.g. visual features, cognitive signals, and other task-relevant or irrelevant information; sometimes called the representational geometry(*36*, *37*)) places important constraints on the role a neural population plays in a particular computation. Area V4 is particularly well-suited to the kinds of decisions and flexibility we studied here because it contains representations of the visual features we varied(*28*, *38*), it is modulated by processes like feature attention, task switching, motor planning, attention, and surround modulation(*5*, *22*, *31*), and our core tasks required judgments about stationary visual stimuli.

Tasks that require subjects to integrate across time or space to perform sequences of decisions or actions might rely more on areas with more complex dynamics(*39–41*) and fundamentally different formatting of task-relevant information.

### Origin of reformatting signals

While the hypothesis tested here predicts how response modulations transform neural population responses, it is agnostic on the origins of those modulations. In our RNN modeling, signals about the task (for example, the location and size of the response arc) are fed into the network explicitly (Figure S8 and S9). These inputs rotate population representation such that both the stimulus curvature and the appropriate eye movement can be decoded linearly. In the brain, such explicit input to V4 is a possible mechanism could originate from a combination of sources, including 1) long-range feedforward connections between early visual areas and V4 across hemifields (since the arc is presented in the upper hemifield)(*42*), 2) recurrent surround modulation within V4(*43–45*), 3) direct or indirect feedback from either visual areas with larger receptive fields (like inferotemporal cortex) or from association areas that remap visual inputs to guide eye movements (e.g. the lateral interparietal area or area 7a)(*46*, *47*).

In the future, causal experiments like those used to establish the origins of modulations related to spatial attention(*48–50*) will be important for establishing the origins of signals related to other forms of cognitive flexibility. Similarly, theoretical work investigating the properties of control signals related to these functions will be important for establishing the feasibility of unified mechanisms for a broader range of flexible behaviors.

### Implications and applications of gain-based reformatting

Together, our results establish the feasibility of a computational mechanism for behavioral flexibility that is rapid, low-cost, robust, and consistent with previous observations about how cognition affects single neurons. We demonstrated that well-known, modest modulations of neural activity can dramatically change the information that is communicated via stable mechanisms. This mechanism may comprise a substrate for the remarkable flexibility inherent to many species and systems. It could provide inspiration for future neuroscientific studies in many species and systems, translational efforts to enhance and repair cognitive flexibility in human patients, and efforts to create biologically inspired artificial systems that mimic the remarkable flexibility of the human brain.

## Acknowledgements

We are grateful to K. McKracken for providing technical assistance, to Douglas Ruff, Mehrdad Jazayeri, Ramon Nogueira, and John Maunsell for comments on an earlier version of this manuscript and for helpful comments and suggestions regarding data analysis. This work is supported by Eric and Wendy Schmidt AI in Science Postdoctoral Fellowship (to R.S.), National Eye Institute of the National Institutes of Health (award K99 EY035362 to R.S.), the Simons Foundation (Simons Collaboration on the Global Brain award 542961SPI to M.R.C), and the National Eye Institute of the National Institutes of Health (awards R01EY022930, R01EY034723, and RF1NS121913 to M.R.C).

## Author Contributions

RS: Conceptualization, Methodology, Software, Formal Analysis, Investigation, Data curation, Writing – Original Draft, Writing – Review & Editing, Visualization, Funding acquisition; MCC: Investigation, Data curation, Writing – Review & Editing; MRC: Conceptualization, Methodology, Formal Analysis, Writing – Original Draft, Writing – Review & Editing, Supervision, Project Administration, Funding acquisition

## Conflict of Interest

The authors declare no competing financial interests.

## Data and Code Availability

The data and code that perform the analyses and generate the figures in this study have been deposited in a public GitHub repository https://github.com/ramanujansrinath/flexigain. Further information and requests for data or custom MATLAB code should be directed to and will be fulfilled by the corresponding author, Ramanujan Srinath.

## Methods

### Experimental Subject Details

#### Continuous curvature report experiments

Two adult male rhesus monkeys (Macaca mulatta, 8 and 9 kg) were implanted with a titanium head post before behavioral training. After training, we chronically implanted an 8×12 microelectrode array in each of cortical areas V1 and V4 of the right hemisphere. (Blackrock Neurotech, Salt Lake City, UT). The arrays were connected to a percutaneous pedestal which allowed for recording. Each electrode within the array was 1mm long, and the distance between adjacent electrodes was 400 μm. Areas V1 and V4 were identified in by visualizing the sulci during array implantation and using stereotactic coordinates. All animal procedures were approved by the Institutional Animal Care and Use Committees of the University of Pittsburgh and Carnegie Mellon University where the electrophysiological and psychophysical data were collected.

#### Shape-color experiments

One adult male rhesus monkey (Macaca mulatta, 10 kg) was similarly implanted with a titanium head post before behavioral training. An 8×8 multielectrode array was implanted in V4 in the left hemisphere. All animal procedures were approved by the Institutional Animal Care and Use Committees of the University of Chicago where the electrophysiological and psychophysical data were collected.

### Experimental Methods

#### Experimentation Apparatus

We presented visual stimuli on a 24” ViewPixx monitor (calibrated to linearize luminance using X-Rite calibrator; 1920 × 1080 pixels; 120 Hz refresh rate) placed 56-60 cm from the monkey. The onset of stimuli was coincident with the onset of a marker on the screen which was captured by a photodiode and recorded (at 30KHz) to synchronize the display with data acquisition. We monitored eye position using an infrared eye tracker (EyeLink 1000 Plus; SR Research) and recorded eye position and pupil diameter (at 2Khz). Neuronal responses were acquired (at 30KHz) using a CerePlex E headstage and CerePlex amplifier (Blackrock Neurotech, Salt Lake City, UT). Reward was delivered using a gravity-based solenoid system (Crist Instruments, Hagerstown, MD)/ Behavioral monitoring, reward delivery, and stimulus rendering were controlled by custom MATLAB software and the Psychophysics Toolbox (v.3; (*51*)).

#### Shape stimulus generation

We used a custom generative algorithm to build and render shape stimuli. Details of generation parameters are schematized in Figure S1. Briefly, random closed loop b-splines (with parameterized complexity) were used to create the 2D cross-sectional shape. These shapes were stacked in the 3rd dimension along the curved medial axis while being scaled and rotated (the length, curvature, thickness profile, and helical twist profile were all randomly generated per shape). This generated the vertices of the 3D shape which were connected with edges and smoothed. The global shape parameters (x, y position on the screen, in-plane and out-of-plane rotation, color, and gloss) were then randomly selected and applied. The shape was rendered in perspective view with the rendering camera directly in front and the light source directly above the camera to create the appropriate shading and specularity cues for the 3D shape. The image was saved and displayed on the screen during the experiment. For the shape-color task, the intermediate/homeomorph shapes between an equilateral triangle and its circumcircle were generated using linear interpolation and displayed on the screen in five colors sampled equally between blue to white or gray.

#### Behavioral Task and Training

Curvature Estimation Task: The monkeys were trained to perform the continuous curvature estimation task schematized in Figure 1. The lengths of each epoch during each trial were sampled from uniform distributions with the following ranges. Monkeys fixated a central fixation point (within a 0.5° fixation window) for 200-250 ms before the shape stimulus was displayed for 500-800 ms. A thin, white arc that usually subtended 140° at the center and a radius of about 3.7° of visual angle was then displayed in the upper hemifield between -70° and 70°. In subsets of sessions, the arc was rotated around the center subtending either between -90°—50°, - 70°—70°, -50°—90° (arc length 140°), or -70°—30°, -50°—50°, -30°—70° (arc length 100°). The radius of the arc was also varied between 3.67-7.3° of visual angle in some sessions. Since the medial axis of the shape was a circular arc, the curvature value the monkey had to guess was the inverse of the radius of the circle.

This curvature was linearly mapped to the target arc. After 180-220 ms, the fixation point was turned off which served as the go cue for the monkeys to saccade to the correct portion of the target arc to make a continuous report of the curvature of the stimulus. After the monkeys maintained fixation on the selected arc location for 200 ms, a juice reward was delivered the magnitude of which depended on the angular error in the curvature report. The reward amount varied from 0.2-0.1 cc up to an error of 10% of the arc length. After the reward was or was not delivered, the monkeys were given feedback: the contrast of the target arc was lowered to reveal a 2° bright spot over the correct target location. Monkeys were rewarded with a small amount of juice to saccade to and fixate the correct location. This second saccade served only as an error correction signal and was not analyzed as an additional or corrected choice. A full trial took between 1.4-2s and all trials in which the monkeys successfully completed the trial (rewarded or unrewarded) were analyzed. Trials with breaks in fixation during any task epoch were not analyzed. The reward strategy was consistent across training, only varying in amount across sessions. Details of session and neuron counts are specified in each figure and analysis. Comparisons of behavioral performance across shapes, orientations, colors, and across trained vs novel shapes are highlighted in Figure S2.

Shape-Color Task: The monkeys were trained to select one of two displayed stimuli based on two rules (schematized in Figure 4). (1) If the color of the stimuli was the same, then the monkey selected the more circular stimulus (shape rule). (2) If the shape of the stimuli was the same, then the monkey selected the bluer stimulus (color rule). Stimuli were sampled from a 5×5 grid of stimuli (5 homeomorphic shapes between a circle and a triangle and five colors between white/gray to blue). Shape and color rule trials were interleaved and the comparison between the two displayed stimuli was the only cue for the relevant rule. The monkeys fixated a central fixation point for 150-250 ms before the stimuli were displayed. The fixation point turned off after 200-250 ms which served as a go cue for the monkey. The monkey made a saccade to one of the stimuli and maintained fixation for 100 ms before receiving a reward for selecting the correct answer.

#### Electrophysiological Recordings and Response Epochs

We filtered the recorded activity (bandpass 250-5000Hz) and detected timestamps where the activity crossed a channel-specific threshold (set to 2-3x RMS value). The raw electrical signal, waveforms at each crossing, and local field potential activity filtered with 2-200Hz were also saved. The timestamps were chunked by trial start and stop times and aligned to stimulus onset determined based on the photodiode signals. To allow for the latency of responses in visual cortex, stimulus-evoked firing rate of each channel was calculated between 50-550 ms (curvature task) and 50-200 ms (Curvature-color task). The baseline firing rates were calculated based on the spike counts in the 200 ms time period prior to the onset of the stimulus. Across 124 sessions, stimulus evoked firing rates ranged from 0-403 (mean 107) spikes/s for V1 units and 0-358 (mean 109) spikes/s for V4 units calculated between 50-550ms after stimulus onset. During the arc presentation duration (0-200ms after arc onset) firing rates ranged from 0-349 (mean 88) spikes/s for V1 units and 0-344 (mean 91) spikes/s for V4 units. Mean baseline firing rates were 59 (V1) and 68 (V4) spikes/s. Analyses comparing single and multiunits in previous work(*52*, *53*), did not find systematic differences between single neurons and multiunits for the types of population analyses presented here. For that reason, we did not sort spikes, and the term ‘unit’ refers to the multiunit activity at each recording site.

#### Unit and trial exclusion

In both tasks, we opted for a generous criterion for units and trials. We included units for analysis if the average stimulus-evoked firing rate across all trials in an experimental session was 1.1x the average baseline firing rate (calculated before the stimulus onset while the monkey was maintaining fixation). Only trials in which the monkeys maintained fixation throughout the stimulus period and made a successful selection were included.

#### Receptive Field (RF) Mapping

The receptive fields of the populations of V1 and V4 neurons were mapped using the same procedure. The monkey fixated a central spot while 2D and 3D object, and texture images were flashed in a grid of locations in the relevant visual quadrant for 200-250ms with a 200-250ms inter-stimulus interval. Monkeys were rewarded to maintain fixation for 6-8 flashes. The images were sampled from a diverse set of random 2D closed-loop b-spline objects, 3D objects generated using the same procedure as for the curvature task (but not used in the curvature task), 2D silhouettes of those objects, and natural texture patches. The images were flashed in at least two sizes to ensure sufficient overlap between locations. For each unit, we calculated a stimulus-triggered average image and an ellipse that fit the RF envelopes. The centers and average sizes of the V1 and V4 receptive fields are shown in Figures S3 and S10. We did not exclude units on the basis of RF location or size.

### Statistical Analysis and Quantification

#### Behavioral performance and psychometric curves

Curvature task: The monkeys’ choice was taken as the angular location on the arc where the monkey entered the arc-shaped window and maintained fixation. The angular location was linearly mapped onto curvature. Psychometric curves in Figure 1 and Figure S2 were plotted by binning actual curvatures across sessions and calculating the mean and standard error of the mean of monkey choices.

Curvature-color task: Since the monkey was rewarded for choosing the more circular (curvature task) or bluer (color task) of the two presented stimuli, each shape stimulus could be compared to four other stimuli in either task. The psychometric curve was calculated as the proportion of trials in which the monkey selected the stimulus in the receptive field as a function of the difference between the stimuli (ranging from -4 to +4, excluding 0).

#### Simulations of decoding performance with fixed readout

To test if typical gain modulations could transform neural population activity to align with a fixed readout axis, we simulated tuning functions to either one (Figure 2-3) or two features (Figure 4 and Figure S11). We simulated 100 neurons with Gaussian tuning functions (results are similar for different numbers of neurons provided n >> 2 to ensure a sufficiently large response space compared to the feature manifold). The tuning width, amplitude, and preferred feature value were randomized. We then calculated the response of these neurons to 2000 stimuli that varied in both features. We selected a random readout axis and then searched for neuron-specific gains such that (1) the neural responses could be transformed to arbitrary sections of the fixed readout axis to resemble flexible alignment to an output axis in the curvature task based on the arc condition, or (2) either of the two feature representations could be aligned to the fixed readout axis to resemble the flexible color- or shape-based decisions in the shape-color task. The gains were strictly constrained to small values such that the gain modulation was similar to previously reported modulations associated with surround modulation or feature attention ((*22*, *43*, *45*); between -20 to 20% difference in firing rates across task conditions), and, like modulations associated with attention, depended on the feature selectivity of the neuron (Figure 4B). After the gains were found, we determined whether the modulated responses for the two arc conditions (Figure 3B) or the two feature conditions (Figure 4B right) aligned with the output axis. To determine whether a similar transformation was a simple consequence of the distributions and feature-dependence of the gain changes, we shuffled the gains such that selectivity-dependence was preserved and recalculated the alignment with the output axis.

#### Curvature selectivity

We calculated the shape-specific curvature selectivity of each unit by first calculating the presentation-averaged curvature response function (Figure S3D) and then calculating the selectivity as (rMax-rMin)/(rMax+rMin) where rMax and rMin were the maximum and minimum response (Figure S3E-F). We calculated this using the stimulus-evoked response (50-500 ms after stimulus onset) or using the response during the arc presentation period (while the monkey was still fixating; 0-200 ms after arc onset). For each shape, the curvature values that corresponded to the maximum and minimum responses were maintained while calculating the selectivity during the arc presentation period. So the selectivity during the stimulus period has a range of 0-1 but not during the arc period.

Principal components analysis, curvature representation, and shape-specific decoding We used Principal components analysis (PCA) primarily for visualizing the population activity and demonstrating a smooth representation of the relevant continuous feature, curvature. We also typically reduced the dimensionality of the population responses to 20-25 before performing decoding or cross-decoding analyses (details below) to prevent rank deficiency. In the population response visualizations (Figure S4, Figure 4G), we plotted each stimulus presentation, where feasible, and the presentation-average responses and color-coded it based on the feature value (curvature, color, or shape). We also fit a 3D polynomial or linear space curve to highlight the relevant variation axis.

For shape-specific curvature decoding (Figure S4), we linearly decoded the curvature values of the presented stimulus from V1 and V4 responses. For display, we binned curvature values and plotted the mean and standard error of the mean (SEM) of decoded curvature values. Unless otherwise mentioned, all decoding analyses were performed with leave-one-out cross-validation (10- and 20-fold cross-validation were also performed with qualitatively similar results; not shown). We typically used the correlation (r) between the decoded and actual feature values as an indicator of decoding performance. We obtained qualitatively similar results using r^2^, mean squared error, and other measures of similarity.

#### Cross-shape curvature decoding

To evaluate whether two shape-specific curvature axes were typically aligned, we split each shape response into equal numbers of trials. We trained a linear decoder using the shape responses for one shape and predicted either the held-out trials of the same shape (Figure 2C) or the other shape (Figure 2D) controlling for trial numbers. We did this in both directions for all pairs of shapes yielding 844 shape pairs across 83 recording sessions. We calculated the mean squared error of self- and cross-decoding and plotted them for shape pairs that differed in overall shape, orientation only, or color only (Figure 2E). For each shape pair, we also calculated the correlation between the choices for the same curvature values and plotted those correlations across the same sessions (Figure S5).

#### Shape-general curvature decoding and behavior correlations

To decode curvature using a shape-general strategy, we trained a linear decoder on responses across different shape stimuli within the same recording session. Since the curvature axes are demonstrably non-aligned, we selected a number of trials to train the shape-general decoder such that the shape-specific and shape-general decoders both have equal curvature decoding performance. To do this, we gradually increased the number of training trials by randomly sampling trials across shapes (across 100 folds) and then comparing curvature decoding performance with the specific decoder of that shape. It took about 40% more trials to train a shape-general decoder to match the curvature decoding performance of the shape-specific decoder. We then compared the decoded curvature values from both decoders to the monkey’s curvature estimation behavior. This yielded two correlation values which are plotted in Figure 2F. We also compared the curvature decoding performance of the shape-general decoder with the monkey’s average curvature estimation error (Figure 2G), and the curvature estimation errors of pairs of shapes within a recording session (Figure 2H). All comparisons were tested for significance using Wilcoxon Signed Rank test.

#### Saccade decoding and intercept analysis

We calculated the spike rate of each unit in V1 and V4 during the arc presentation duration (0-200 ms after arc onset) and linearly decoded (using leave-one-out cross-validation) the actual saccade locations on the arc in arc degrees. We then compared the actual saccade and the decoded saccade split between arc conditions. (Note that we did not fit separate decoders for each arc condition; we instead used an arc-general saccade decoding strategy.) We tested the hypothesis that V1 and V4 responses would transform to encode the curvature and the saccade simultaneously by realigning/rotating to have an explicable projection on the saccade axis. The null hypothesis would be that V1 and V4 responses would continue to only represent the stimulus in their RFs. If the null hypothesis was true, the relationship between the actual and decoded saccades would mimic the relationship between the actual saccade and the stimulus curvature and therefore be arc dependent. We tested this by calculating the intercept of this relationship across different arc conditions per stimulus shape both before and after arc onset. In many cases, we tested several arc conditions (variations in both arc length and position) for the same shape and several shapes in the same experiment therefore yielding curvatures x shapes x arc location x arc position trial conditions.

#### Recurrent neural network (RNN) modeling of saccade outcomes

We trained an RNN using PsychRNN(*54*) to understand the possible computational mechanisms of the transformation between stimulus and saccade representations. We input the fixation as a one-hot variable, the curvature as a continuous variable between 0 and 1, and the arc condition as two variables that represented the arc location and length (Figure S8A). Three arc angular positions and two arc lengths were tested. The RNN was trained to calculate the correct saccade for the input variable combination. The timing of the different inputs was randomized and sequenced to replicate the monkey task (schematized in Figure S8B). The RNN had 50 hidden units and was trained on noisy, random inputs. The network learned to do the task (compare Figure S8D and S8F) by first representing the curvature during the stimulus epoch and, after arc onset, rotating the representation to align with a saccade axis such that both the curvature and the saccade can be read out.

#### Decoding curvature from convolutional network (CNN) activations

Since the veridical curvature was supplied as input to the RNN above, we next tested whether a pre-trained convolutional neural network (VGG-16) could be used as a front-end to extract the 3D curvature from image inputs. To do that, we rescaled the input images to the appropriate size and the shape stimulus in the image to the size of the RF of the pooling layer neurons in layer 4 of VGG-16. We generated either 1000 fully random shapes with random curvatures (image set 1), or 50 unique shapes with 20 curvatures each (image set 2). We trained a linear decoder using the activations of the central units (since the stimulus image was scaled and placed in the middle of the image) and tested it on a testing set of 5 shapes with 20 curvatures each. We reduced the dimensionality of the response space to 256 dimensions to prevent rank deficiency. The curvatures of the test set were successfully decoded by shape-general decoders trained on either of the image sets (Figure S9A).

#### RNN saccade modeling using CNN inputs

We then supplied an RNN with the dimensionality-reduced CNN inputs and trained it using the same method described above. We used a larger network (512 hidden units) to compensate for the high dimensional input but otherwise kept the training regimen and trial timing the same as above. We trained separate RNNs of the two image sets and both were able to generate the correct saccades (Figure S9B; network behavior) by first representing the curvature on the stimulus axis and then rotating the representation in an arc-dependent manner to the saccade axis (Figure S9B; network state).

#### Shape-color tuning, selectivity, and task gain

We calculated the stimulus-evoked spike rate of each V4 neuron 50-200 ms of stimulus onset to account for spike latency to V4. We plotted the trial-averaged responses for each of the 25 stimuli to find that neurons across the array had a variety of tuning peaks and widths (Figure S10C). We quantified the feature selectivity of each neuron as (rMaxColor-rMaxShape)/(rMaxColor+rMaxShape) where rMaxColor and rMaxShape were the maximum responses along the color and shape dimensions across both task conditions. We also calculated the task gain for each neuron as the same modulation ratio except where rMaxColor (or rMaxShape) was the maximum response across stimuli during the color (shape) task. The selectivity and gain were calculated for 23 recording sessions (41-48 neurons each, median 45) yielding 1033 points.

#### Shape-color-choice representation and decoding

We first visualized the shape and color responses of the V4 neurons by either reducing the dimensionality using PCA and plotting the projection of each trial on the first three principal components or by identifying the shape and color decoding axes using linear regression and using QR decomposition to find orthogonal bases (Figure 4G). To find the feature and choice decoding axes, we used the same decoding procedure as the cross-decoding results described above. Briefly, we randomly split the trials from the shape and color tasks equally into training and testing groups. We used the training group to linearly decode the shape or color of the stimulus and tested it by using the same weights to predict the shape, color, or choice on the testing trials. We trained separate shape and color decoders each for the shape and color task trials and one for both tasks combined (totaling six decoders) and tested each of them on shape, color, and choice decoding (Figure 4H and Figure S12).

## Supplementary Figures

**Figure S1:**
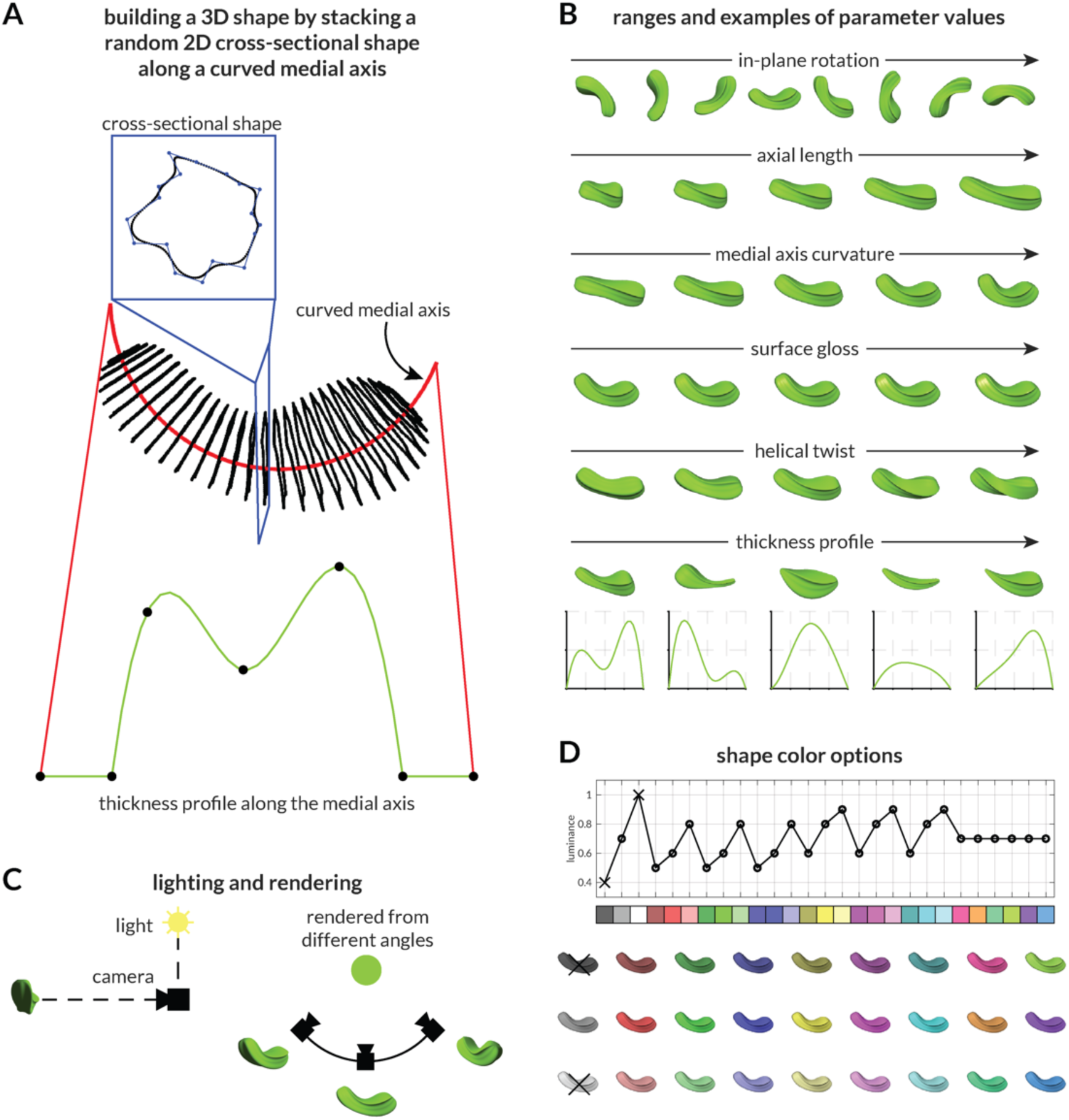
Stimulus generation details and parameters. **A:** To build a random 3D stimulus shape, we first generate a cross-sectional 2D shape by randomizing a closed-loop cubic b-spline with arbitrary complexity (number of control points and randomization). We also generate a curved axis with a randomized length and curvature, a thickness profile along the axis (by randomizing a 5-point Bezier curve), and a helical twist profile along the axis. We then scale the cross-sectional shape by the thickness profile and stack it along the axis. Once the vertices are in place, we join them with edges to create a closed 3D object. We render the object with a randomized color, position, in-plane orientation, and surface specularity/gloss. **B:** Example renders of the same object with changes in different features. Wherever possible, the ranges spanned the parameter space (orientation, color, curvature, twist, etc.) and were otherwise chosen manually (length, gloss, thickness profile). **C:** The lighting and camera were held in the same positions across all images – the camera was directly in front of the object and the light was an omnidirectional source above the camera. On the right, for demonstration, the object is shown being rendered along three camera positions. **D:** The shape colors were chosen from 25 random values which were generated by permuting three absolute R, G, and B values – 0.4, 0.7, 0.9. The swatches and average luminance values thus generated are shown at the top and the example stimulus rendered in those values are shown below.

**Figure S2:**
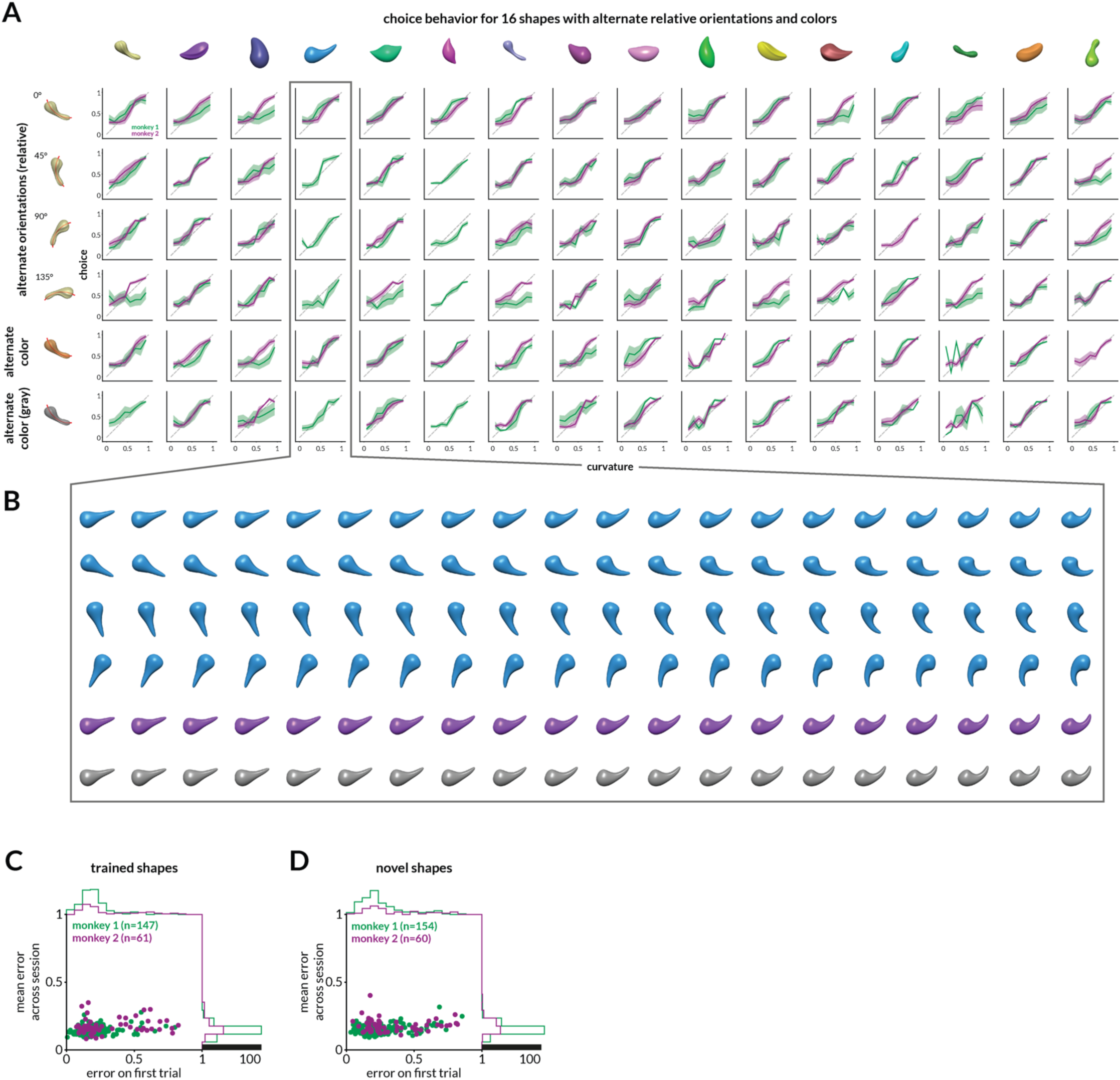
Curvature estimation behavior across all tested shapes. **A.** Curvature estimation behavior shown as psychometric curves for the two monkeys (blue and orange lines, respectively) for all unique shapes tested (the medial axis of each shape, shown in red in the icons, was not presented to the monkeys and is shown for illustration). Each panel shows the comparison between the actual curvature and reported curvature in solid lines (in nine bins across the curvature range). Shading indicates SEM within each bin. For each shape tested, 10 or 20 curvature values are presented as shown in the icons in the exploded view (in B) for shape 4 as an example. Each column represents a unique shape. The first four rows represent orientation variations as shown in the icons on the left and in the exploded view. The last two rows similarly represent color variations. Both monkeys tended to underestimate high curvature values and overestimate low curvature values. This might be a result of the bounded nature of the curvature report and/or anisotropies in curvature representations across the range of curvatures tested. **B.** An exploded view showing all stimuli tested for a single shape. In the text, we use the term ‘shape’ to refer to a full set of curvatures with no other parameter changes. Across shapes, the orientation only, the color only, or the entire shape (including many other parameters) could change. **C.** Comparison of absolute behavioral error on the first presentation of a shape (one of 10 shapes used during behavioral training) during a recording session and the mean absolute error across the whole session. For both monkeys, the distribution of errors (marginal histograms on the right and top) tightens across the session and settles at a median value of ∼ 0.12. **D.** Same as C but for shapes never seen before during recording or training. The distribution of errors for both monkeys is similar to the one seen in C for the first trial and across the session.

**Figure S3:**
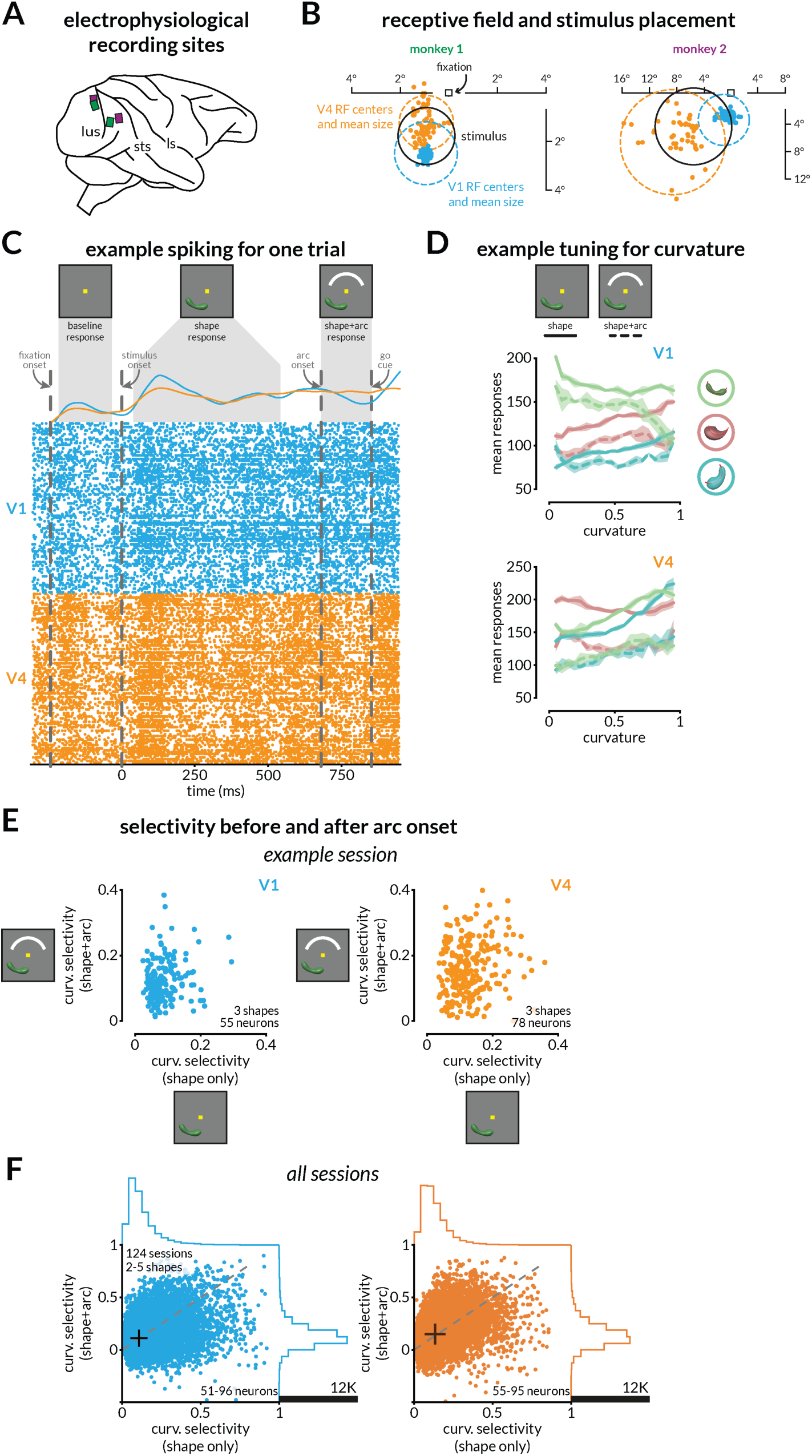
Recording locations, receptive fields, example responses to curved shapes, and neuron-wise curvature selectivity across shapes. **A.** Locations of the multielectrode array implantations for the two monkeys. **B.** Locations (dots) and mean size (dotted circle) of V1 and V4 receptive fields. The black circle indicates the stimulus location for an example session. **C.** Raster plot and peri-stimulus time histogram for a single example trial. Each row corresponds to a V1 or V4 multiunit site. The baseline, stimulus, and arc periods (during which spikes are counted toward the response) are labeled in gray. **D.** Trial-averaged responses for an example site in V1 and in V4 for three shapes as a function of curvature. Solid/dotted lines indicate responses during the stimulus/arc period, and shading indicates SEM. **E.** Comparison of selectivity (defined as how much curvature modulates responses of a single shape for one unit; see Methods) during stimulus and arc periods for all shapes and each V1 and V4 site recorded in an example session. **F.** The curvature selectivity shown for all shapes tested for all neurons across all sessions. The selectivity for curvature does not change systematically after the onset of the arc in either visual area. This was calculated for each shape (2-5; mean 3.55), for each recording site (51-96 for V1, 55-95 for V4), and for each of 124 sessions, totaling 34827 points for V1 and 35537 points for V4. Curvatures that elicited the maximum and minimum firing during the stimulus epoch were kept consistent while calculating selectivity during the arc period. Mean selectivity is shown in black + symbols; V1 selectivity changes from 0.13 to 0.142, and V4 selectivity changes from 0.134 to 0.156 after arc onset on average. Marginal distributions are plotted on the top and right of both panels. Gray line indicates the unity line.

**Figure S4:**
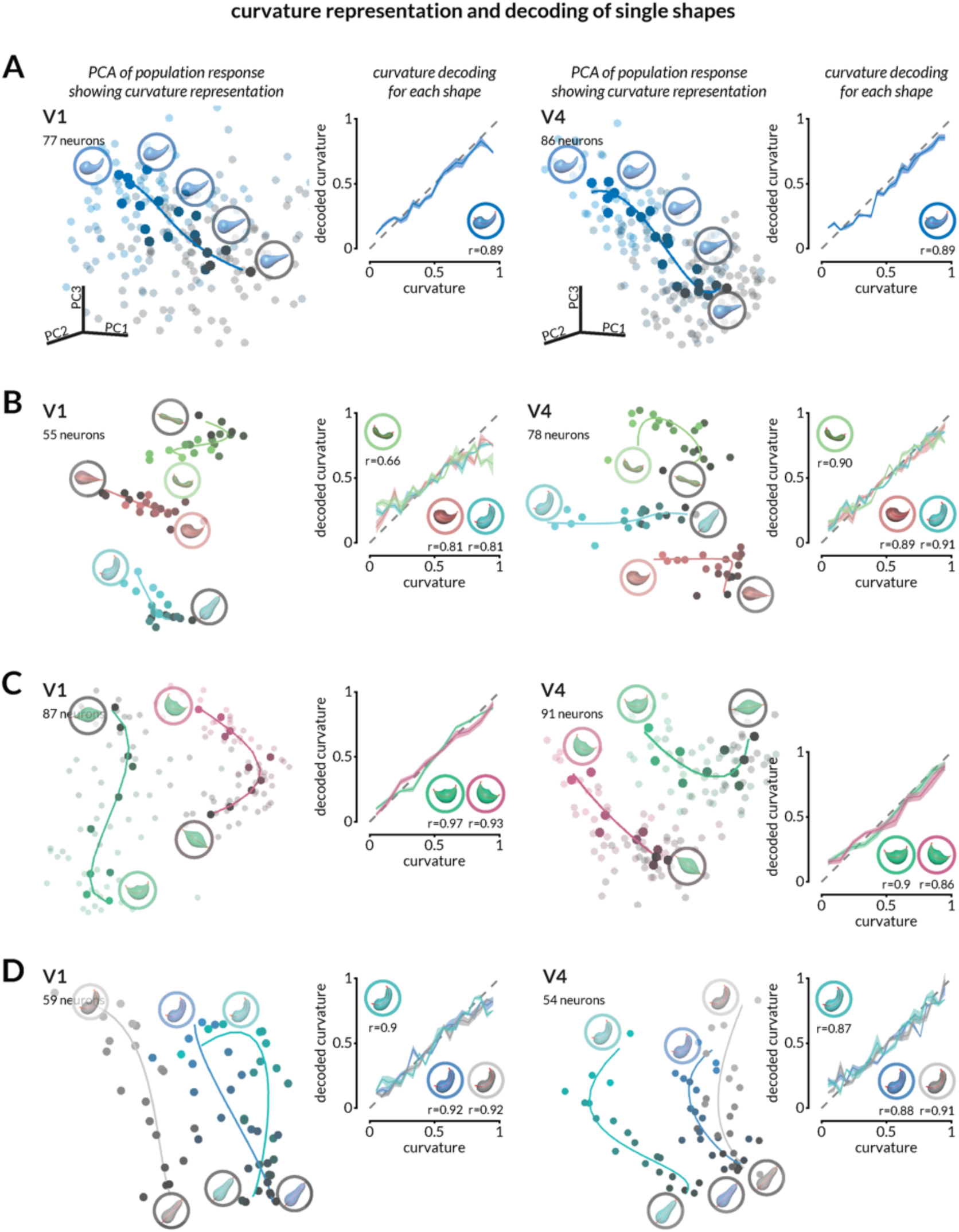
Curvature is robustly encoded in V1 and V4 population activity. (see Figure 1 for choice behavior during the same sessions) **A.** (left) V1 stimulus-evoked population response from 77 neurons recorded during an experimental session projected onto the first three principal components of the population space where each dimension represents the response of one neuron. The principal component analysis (PCA) is for visualization rather than formal analysis. Each dot represents population responses during a single stimulus presentation, and the dot luminance (black-to-blue gradient) represents the curvature of the stimulus (also shown as superimposed icons for a subset of the presented curvatures). The dimmer dots represent all presentations and brighter dots are trial-averaged responses of stimuli with unique curvature. The solid curve represents a polynomial fit along the curvature representation. After training a linear decoder to predict curvature using neural responses, we binned trials by curvature and plotted their average decoded curvature and SEM (middle left); decoding performance was calculated as the correlation between the actual and decoded curvature (here, r=0.89). The same analysis is shown for a simultaneously recorded V4 population of 86 neurons on the right. **B.** Same as A, for three unique shapes shown on randomly interleaved trials in the same session. The decoding performance for each shape is shown under the shape icons in the relevant plots. **C.** Same as A, for the same shape presented in two different orientations. **D.** Same as A, for the same shape presented in three different colors.

**Figure S5:**
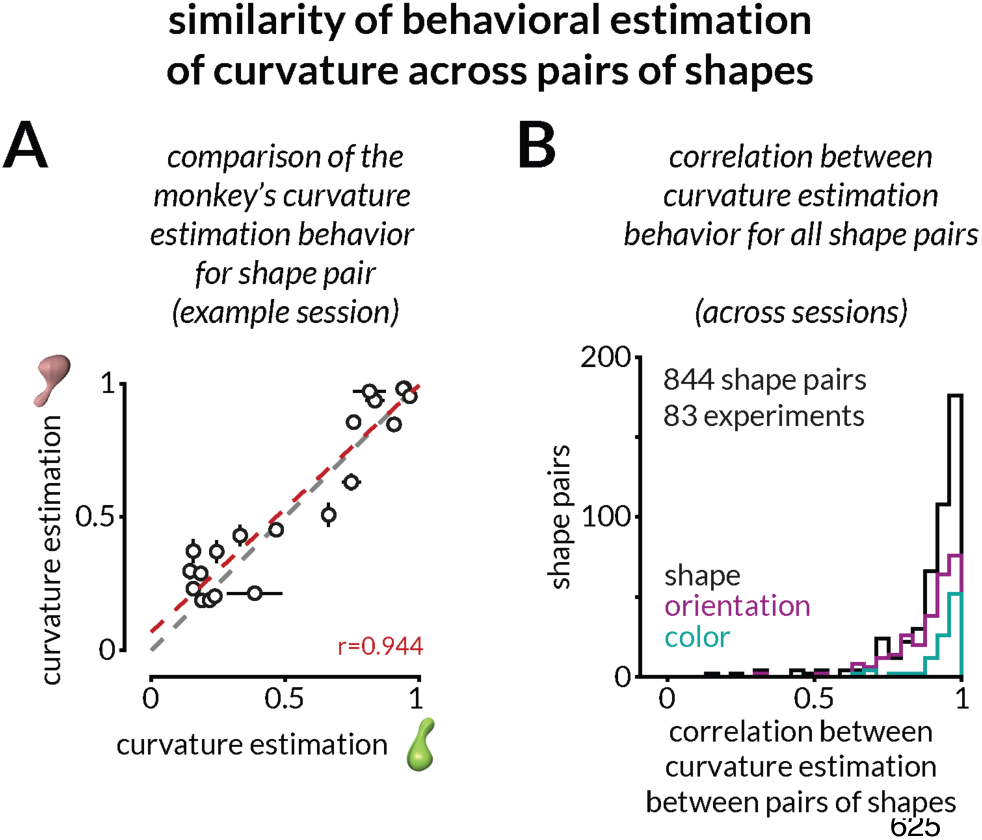
Quantifying the similarity in curvature estimation behavior across pairs of shapes. **A.** While the representations of curvature in visual cortex responses are not typically aligned, the monkey’s curvature estimation behavior is similar across shapes in the same session. Each dot is a unique curvature value, and horizontal and vertical error bars indicate SEM for the two shapes shown as icons. All behavioral biases are consistent across these shapes (correlation coefficient, r=0.944). Compare with curvature decoding in V1 and V4 for the same session in Figure 2. **B.** Monkeys’ curvature estimation behavior for any pair of shapes (unique, orientation-change, and color-change) is consistently correlated across sessions (mean 0.89, SEM 0.01, 844 shape pairs).

**Figure S6:**
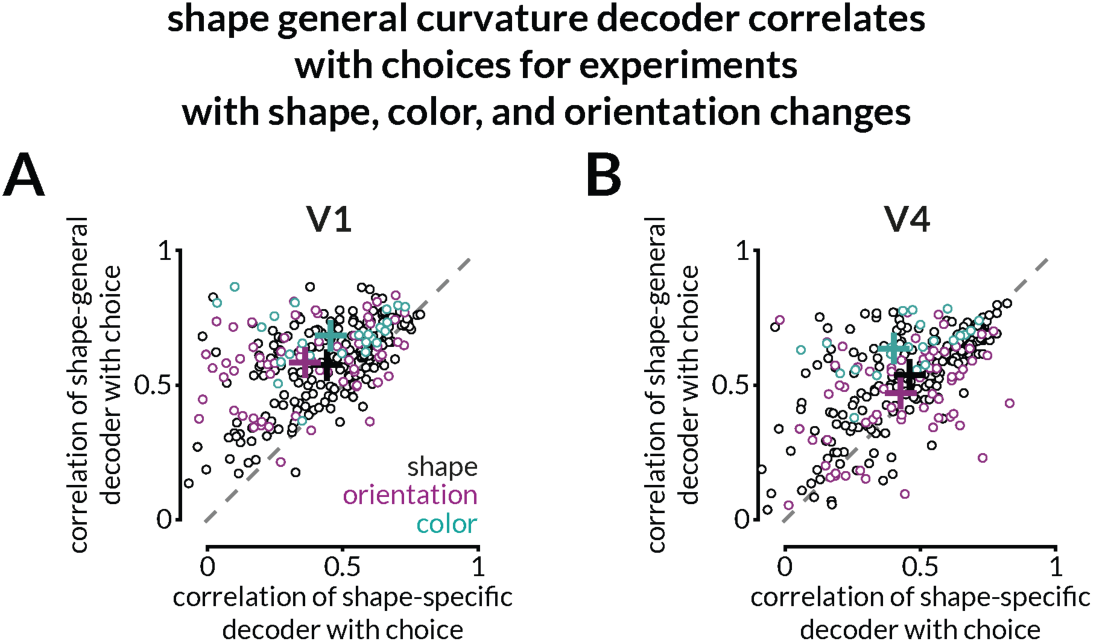
Shape-general decoders trained on responses to shapes differing in color only, orientation only, or overall shape are correlated with choices. Curvature decoded using shape-general or shape-specific decoders (calculated in Figure 2D) split by sessions where only color, only orientation, or overall shape were varied across trials (same splits as in Figure 2F). Curvature decoded with shape-general decoders is more correlated with the monkey’s choices than curvature decoded with shape-specific decoders for all training set variations (Wilcoxon signed rank test; p<0.001).

**Figure S7:**
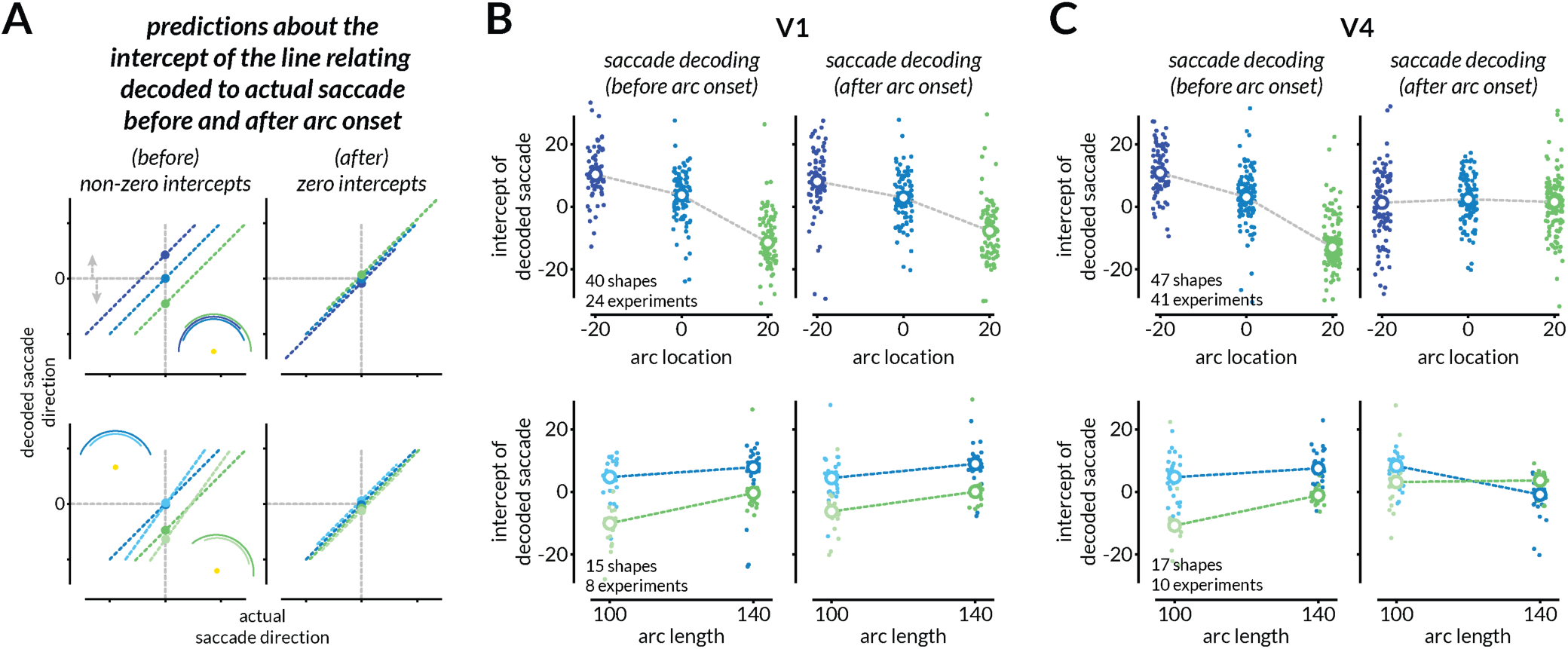
Analyzing intercepts of the linear relationship between the actual and decoded saccade reveals that V4, but not V1 responses reformat to encode the upcoming saccade. **A.** We quantify the extent to which each population reflects the direction of the upcoming eye movement by evaluating the intercept of the line relating the decoded to the actual saccade. If the population does not reflect the direction of the upcoming eye movement, the intercepts will erroneously depend on the arc location (top left). In this scenario, rightward arc shifts will produce a negative intercept (green) and leftward shifts will produce a positive intercept (dark blue). When the arc length is varied, the intercepts will be zeros for centrally presented arcs (blues) but will be different negative values for rightwards shifted arcs (greens) (bottom left). If the population responses reflect the direction of the upcoming eye movement, all intercepts will be zero (top right and bottom right.). **B.** Across experiments, V1 responses reflect the curvature judgment, not the direction of the upcoming eye movement. The intercept of the line relating the decoded and actual saccade direction depends on the arc location both before (left) and after the onset of the arc (right). Each point represents one combination of shape and arc condition. **C.** Across experiments, V4 responses reflect the direction of the upcoming eye movement. Conventions as in B. Before the arc onset, the results are qualitatively similar to V1. Unlike in V1, the saccade direction can be decoded from V4 responses after the arc onset (as indicated by zero intercepts across arc conditions; right panels).

**Figure S8:**
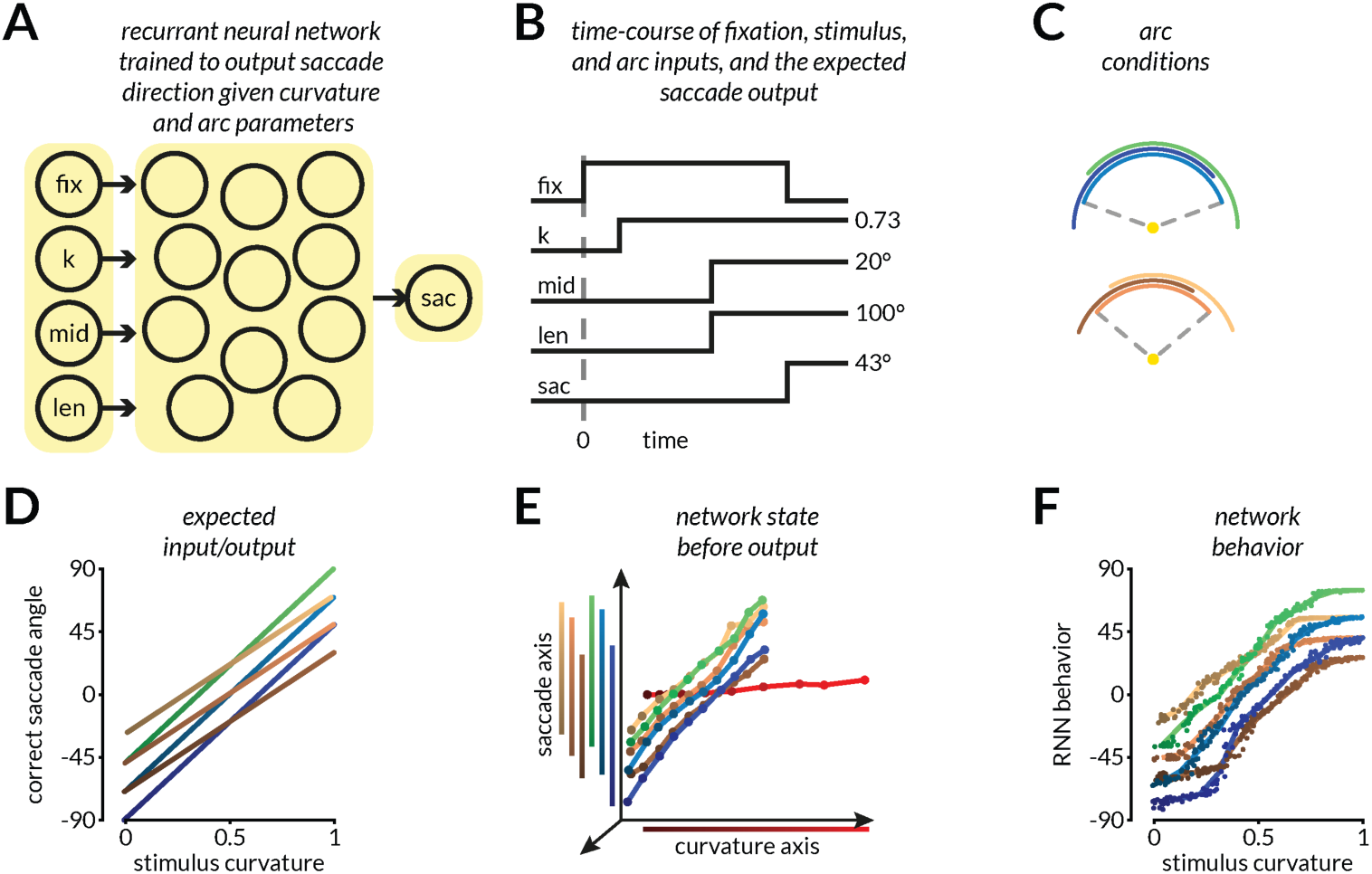
Recurrent neural networks (RNNs) can be trained to represent continuous variables and reformat those representations based on stimulus-extrinsic inputs in ways that could produce saccade-like outputs. **A.** Schematic of RNN layout. Four time-varying signals that indicate the fixation spot (“fix”), stimulus curvature (“k”), and the two arc conditions – arc location (“mid”) and arc length (“len”) – are input to the network. The network is trained to produce saccade outputs (“sac”) (shown in D) appropriate for each arc condition (shown in C). **B.** The time course and values of input and expected output signals. Notably, the fixation, stimulus, and arc conditions are staggered to mimic the timing in the monkey task. The relative onsets are also randomized using the same timing distributions. **C.** Legend for the various arc conditions tested. **D.** The expected outputs for each arc condition as trained using backpropagation. **E.** The network states (as visualized by a low dimensional embedding of the hidden layer activations using PCA and QR decomposition) before (red line) and after (colored lines) arc onset. We found the curvature and saccade axes by finding the linear decoding axes for curvature (before arc onset) and for saccades (after arc onset). Black-to-red dots each correspond to increasing curvatures of the stimulus and the projection of this curvature representation on the curvature axis is shown in the black-to-red line along the x-axis. The network activations after arc onset are projected onto the same dimensions and then used to decode the output/saccade. The projection of network states after arc onset (but before fixation offset) for each arc condition is shown in black-to-color lines along the y-axis. The layout of these projections recapitulates the expected output of the network (also shown in F). Importantly, the network does not lose the ability to read out stimulus curvature while encoding the output before the saccade. **F.** The output of the trained network compared with the stimulus curvature for each arc condition. Compare with D and note intersecting points across arc conditions.

**Figure S9:**
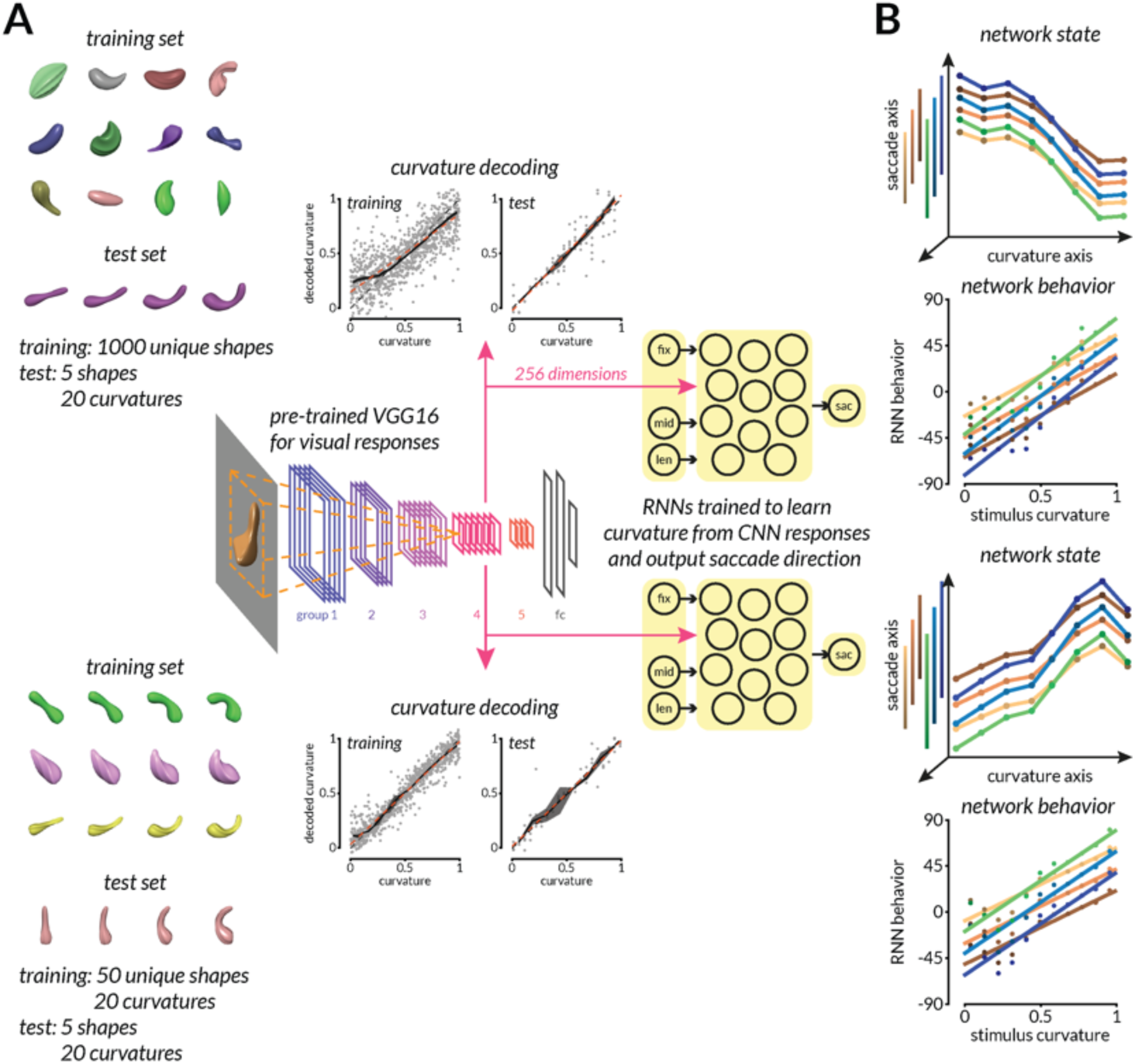
Convolutional Neural Network (CNN) activations can be used to extract visual features and train RNNs to flexibly align representations to behaviorally relevant readout axes. **A.** We extract visual responses from an intermediate layer of a pre-trained convolutional neural network (VGG-16). We use these responses (picked because that layer has been shown to be representationally similar to V4) to train RNNs as in Figure S3. Visual stimuli are created using the same methods as described in Figure 1 and rendered in the central 91 pixels (RF of layer 4 units in VGG16) of the image, and visual responses are extracted. The dimensionality of activations is reduced to 256 to aid in RNN training and linear decoding. Two types of training image sets were created – 1000 unique shapes with randomly varying curvatures (top), or 50 unique shapes each with 20 curvature values (bottom). The testing stimuli in both conditions comprised of five unique shapes with 20 curvature values each. To show that curvature is represented in VGG16 layer 4, we trained a linear decoder using those activations in a leave-one-out cross-validated fashion using training images and tested it on the held-out test images. The same activations, reduced to 256-dimensions, were used to train RNNs instead of the one-dimensional stimulus input in Figure S8. **B.** The trained RNN’s hidden layer activations and output behavior are shown in the two pairs of panels. The top two are for the training set with 1000 unique shapes, bottom two are for the training set with 50 unique shapes. In both cases, the hidden layer represents the stimulus before arc onset, and the representation reformats to align with the saccade-like axis after arc onset. After this realignment, the curvature information is not lost but is still decodable, as in V4 (compare with Figure 3 and Figure S8).

**Figure S10:**
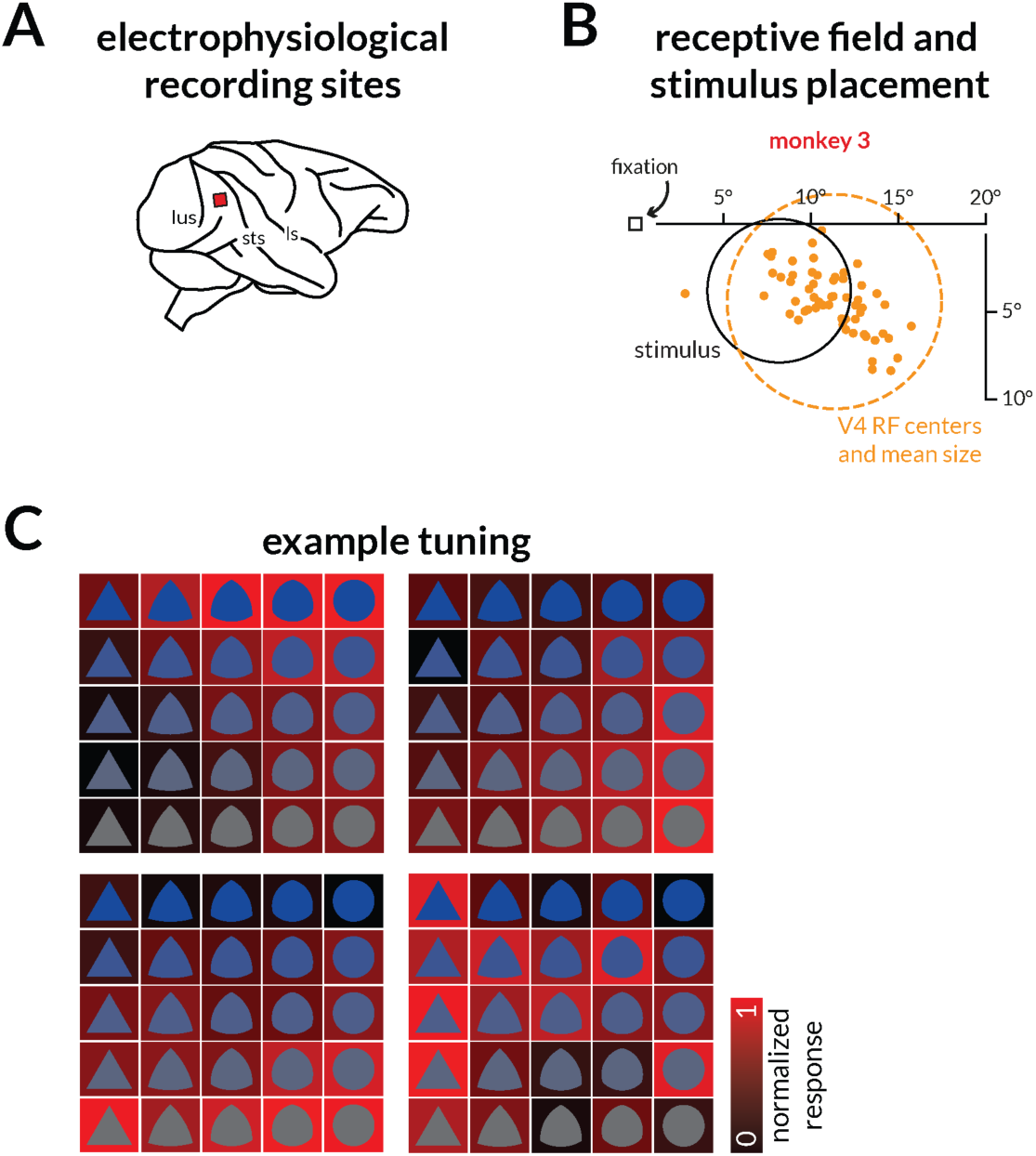
Recording locations, receptive fields, example responses to images used in two-feature 2AFC task. **A.** Locations of the multielectrode array implantations for the third monkey. **B.** Locations (dots) and mean size (dotted circle) of V4 receptive fields. The array location yielded relatively eccentric RFs and the stimulus (black circle) was chosen to overlap with the receptive fields of a majority of the recorded units. **G.** Normalized responses of four example multiunits recorded simultaneously. The black-red saturation of the background of the shape image indicates the normalized firing rate of that neuron. Across the array, the shape and color selectivity of the neurons varied considerably.

**Figure S11:**
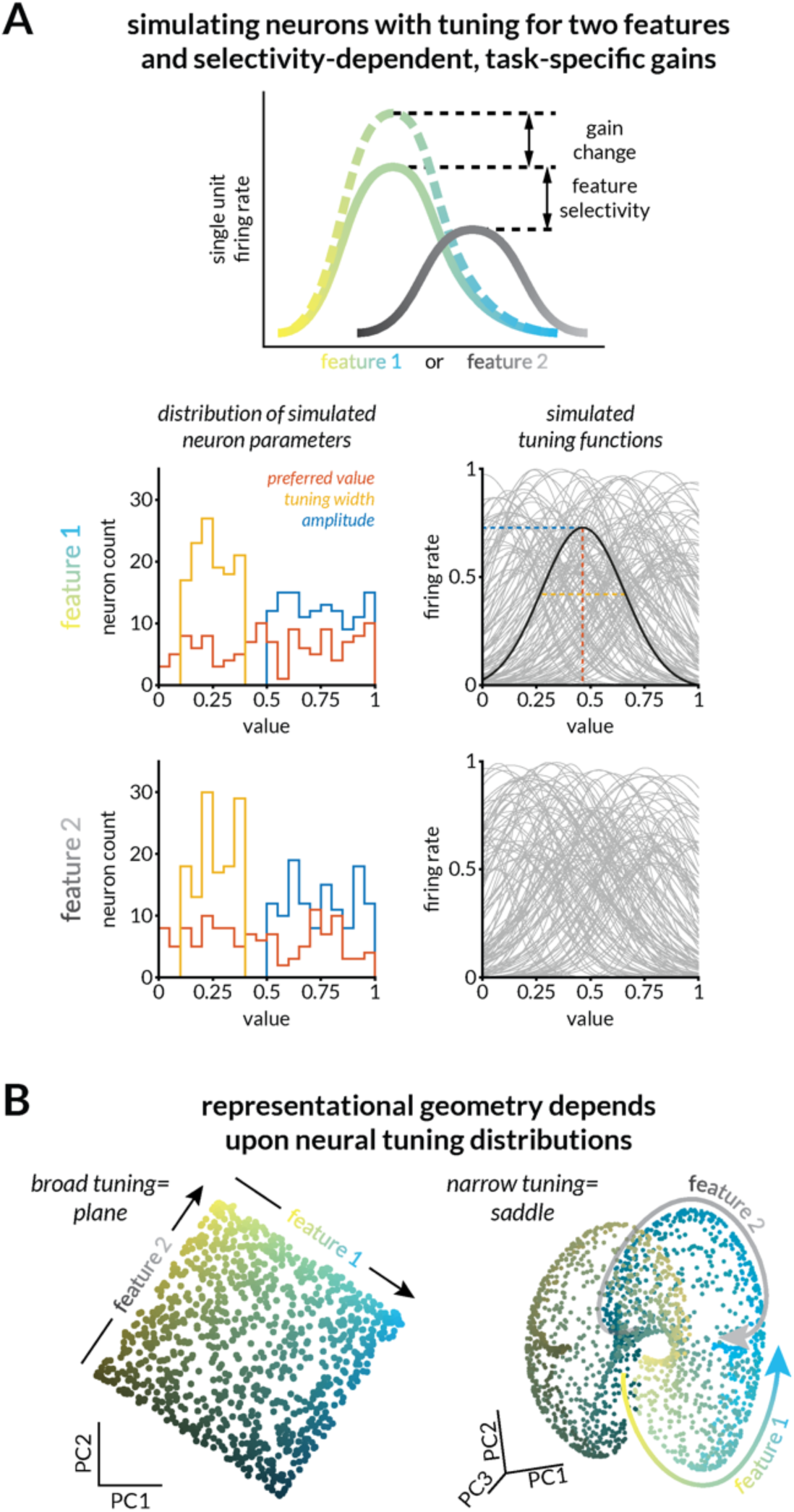
Simulations of tuning functions for two features can result in very different representational geometries. **A.** Gaussian functions for two arbitrary parameters had identical distributions of Gaussian amplitude (0.5-1sp/s), tuning width/standard deviation (0.1-0.4), and preferred value/gaussian mean (0-1). The parameters were drawn from uniform distributions. **B.** Two example population geometries created using different instantiations of the simulation. Here, varying the distribution of tuning widths gives rise to a planar representation or a saddle-shaped representation. In both cases, the bounded nature of the two features causes anisotropies at the edges of the representations causing linear decoders to underestimate higher feature values and overestimate lower values.

**Figure S12:**
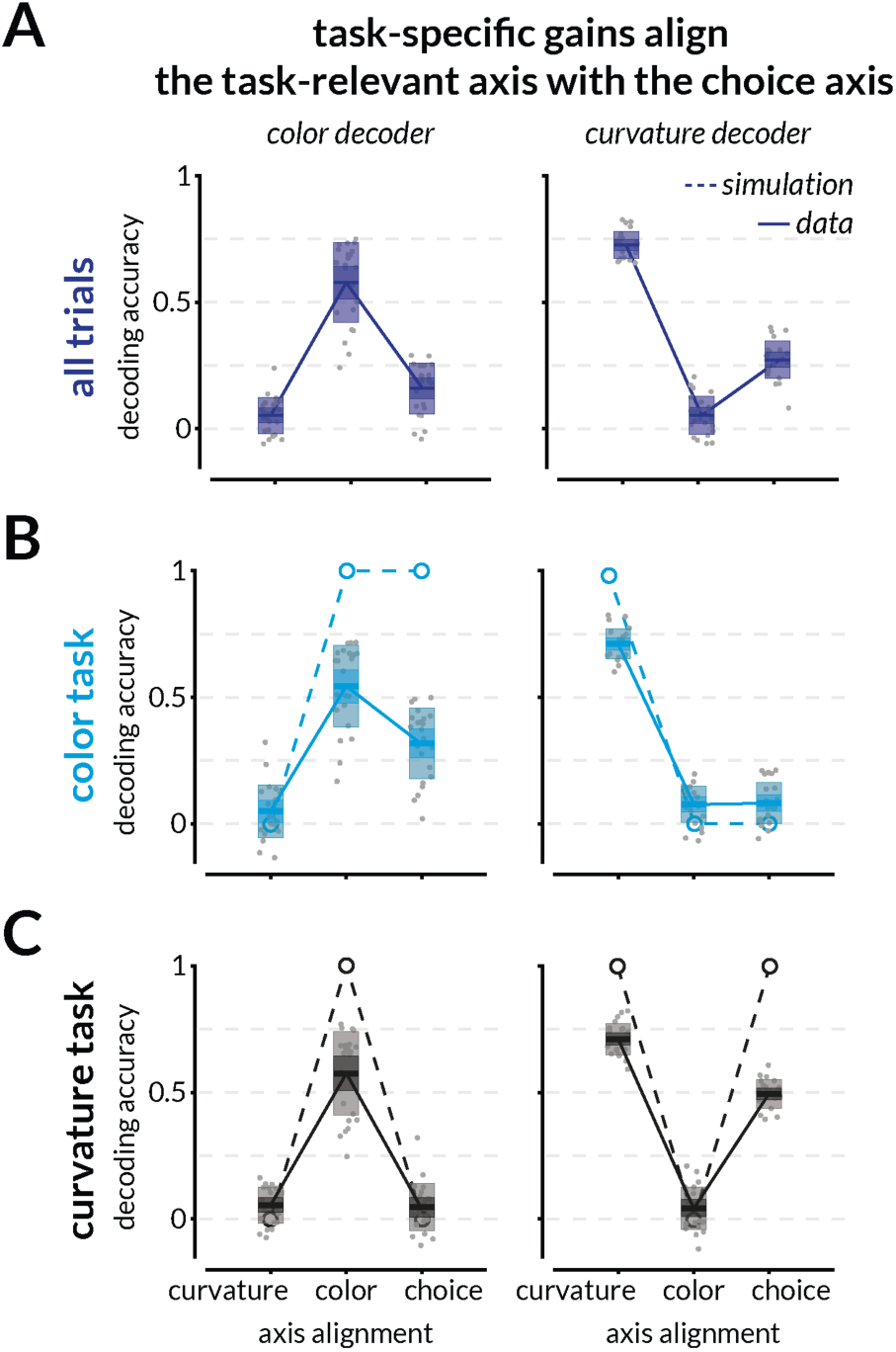
Breakdown of accuracies for the color and curvature decoders trained on either color task or shape task or all trials while decoding color, curvature, or choices. Decoding accuracies (correlation between actual and decoded values) for decoders trained on color (left) and curvature (right) while decoding curvature, color, and choice trained and tested across subsets of (**A**) all trials, (**B**) color task trials, and (**C**) curvature task trials. Points in B and C are identical to those plotted in the scatter diagram in Figure 4H. To illustrate that the shape and color axes are consistent across the two tasks, we decoded curvature, color, and choice across trials from both tasks together (shown in A). Curvature and color decoding accuracies were comparable to the accuracies of those trained using the individual task trials and choice decoding accuracy was approximately halfway between. The open circles/dashed lines depict model predictions.

## Notes

### Competing Interest Statement

The authors have declared no competing interest.

